# Fission yeast Caprin protein is required for efficient heterochromatin establishment

**DOI:** 10.1101/2024.06.19.598224

**Authors:** Haidao Zhang, Ekaterina Kapitonova, Adriana Orrego, Christos Spanos, Joanna Strachan, Elizabeth H. Bayne

## Abstract

Heterochromatin is a key feature of eukaryotic genomes that serves important regulatory and structural roles in regions such as centromeres. In fission yeast, maintenance of existing heterochromatic domains relies on positive feedback loops involving histone methylation and non-coding RNAs. However, requirements for *de novo* establishment of heterochromatin are less well understood. Here, through a cross-based assay we have identified a novel factor influencing the efficiency of heterochromatin establishment. We determine that the previously uncharacterised protein is an ortholog of human Caprin1, an RNA-binding protein linked to stress granule formation. We confirm that the fission yeast ortholog, here named Cpn1, also associates with stress granules, and we uncover evidence of interplay between heterochromatin integrity and ribonucleoprotein (RNP) granule formation, with heterochromatin mutants showing reduced granule formation in the presence of stress, but increased granule formation in the absence of stress. We link this to regulation of non-coding heterochromatic transcripts, since in heterochromatin-deficient cells, absence of Cpn1 leads to hyperaccumulation of centromeric RNAs at centromeres. Together, our findings unveil a novel link between RNP homeostasis and heterochromatin assembly, and implicate Cpn1 and associated factors in facilitating efficient heterochromatin establishment by enabling removal of excess transcripts that would otherwise impair assembly processes.

## Introduction

Heterochromatin is a distinct, condensed form of chromatin that serves important regulatory and structural roles in eukaryotic genomes. It can be divided into two types: facultative heterochromatin contributes to dynamic regulation of gene expression in response to developmental or environmental cues, whereas constitutive heterochromatin is typically found in gene-poor, repeat-rich regions of the genome such as centromeres, where it plays important roles in the maintenance of genome stability including repressing invasive genetic elements and supporting accurate chromosome segregation (1). Heterochromatin is characterised by relatively low levels of histone acetylation, and in most eukaryotes by enrichment for methylation of histone H3 at lysine 9 (H3K9me). This histone methyl mark provides binding sites for chromodomain proteins including heterochromatin protein HP1, which contributes to chromatin compaction and promotes recruitment of further effector proteins involved in transcriptional silencing, chromatin organisation and chromosome segregation (2).

Paradoxically, although heterochromatin is generally associated with transcriptional repression, local transcription is commonly required to support heterochromatin assembly, and RNA processing and surveillance pathways have been shown to play dual roles in co-/post-transcriptional silencing and RNA-based targeting of chromatin modifiers (2–4). In the fission yeast *Schizosaccharomyces pombe*, the RNA interference (RNAi) pathway plays a major role in maintenance of constitutive pericentromeric heterochromatin. Non-coding RNAs derived from pericentromeric repeat sequences are processed by Dicer (Dcr1) to generate short interfering (si)RNAs (5–7). These siRNAs are bound by Argonaute (Ago1) and serve as sequence-specific guides to target the Ago1-containing RITS complex to complementary nascent RNAs (8,9). Via the adaptor protein Stc1 (10,11), RNA-bound RITS mediates recruitment of the Clr4 complex (CLRC) comprising the H3K9 methyltransferase Clr4 along with several other factors including WD repeat protein Rik1, all of which are required for histone methylation (10,12–15). The resulting H3K9 methylation in turn helps to stabilise recruitment of both RNAi and heterochromatin factors via binding of chromodomain-containing proteins including the RITS subunit Chp1, HP1 protein Swi6, and Clr4 (16), such that siRNAs and H3K9 methylation together form a self-reinforcing signal loop for heterochromatin maintenance.

In fission yeast, additional RNA-dependent but RNAi-independent pathways have also been found to contribute to heterochromatin assembly, particularly at domains of facultative heterochromatin. These domains include so-called heterochromatin islands, typically associated with meiotic genes that are repressed during vegetative growth (17,18), as well as additional heterochromatin domains (HOODs) that are detectable at retrotransposons and some developmentally regulated genes in certain conditions, including when the nuclear exosome is inactivated (19). While HOODs are partially RNAi-dependent, heterochromatin islands are RNAi-independent, and both rely to varying degrees on other RNA processing factors including the exosome adapter Mtl1-Red1 core complex MTREC (20,21), the 5′-3ʹ exonuclease Dhp1/Xrn2 (22,23), and the Ccr4-Not deadenylase complex (24–27), for coupled RNA elimination and heterochromatin assembly. Absence of the Ccr4-Not complex has been shown to be associated with increased accumulation of heterochromatic transcripts on chromatin that leads to impaired silencing (24), indicating that while RNA production is required to provide a platform for recruitment of heterochromatin assembly factors, excess accumulation of such transcripts can negatively impact heterochromatin integrity.

A key feature of heterochromatin is its capacity for epigenetic inheritance through reader-writer coupling (28). In fission yeast, high densities of H3K9me3 promote binding of Clr4 and methylation of neighbouring nucleosomes, facilitating propagation of pre-existing heterochromatin (29). This epigenetic maintenance phase can be distinguished from *de novo* heterochromatin establishment, which occurs through two consecutive stochastic steps: initiation at nucleation sites, followed by subsequent spreading to form extended domains (30). Dependent on context, some factors and processes required for *de novo* heterochromatin establishment are redundant for subsequent maintenance, necessitating dedicated strategies for their detection. For example, histone deacetylase (HDAC) activity is required to promote heterochromatin assembly by increasing availability of deacetylated H3K9 residues for methylation as well as suppressing acetylation-dependent histone turnover (31). Deletion of HDACs Sir2 or Clr3 has only modest effects on maintenance of pericentromeric heterochromatin, but strongly impacts *de novo* heterochromatin nucleation and spreading on pericentromeric repeat sequences (32,33). A small number of additional factors have also been identified as being specifically required for heterochromatin establishment with little or no role in subsequent maintenance, including Triman (Tri1), a 3′-5′ exonuclease involved in siRNA biogenesis (34), and the RNA-binding protein Mkt1 that is involved in RNAi-dependent post-transcriptional regulation of heterochromatic transcripts (35). We previously showed that Mkt1, similar to Ccr4-Not complex, is required to prevent excess accumulation of transcripts on chromatin and formation of RNA-DNA hybrids, suggesting that excess RNA may be a particular impediment to the initial conversion of highly transcribed chromatin to heterochromatin.

Here, through an assay involving genetic abolition and re-establishment of heterochromatin, we identify the product of a previously uncharacterised gene, SPAC12G12.07c, as a novel factor required for efficient heterochromatin establishment in *S. pombe*. The SPAC12G12.07c gene product is orthologous to human RNA-binding protein CAPRIN1, and we therefore name it Cpn1. Human CAPRIN1 is involved in stress granule formation, and we confirm that fission yeast Cpn1 both associates with stress granule-related factors, and accumulates in granules during stress. Moreover, we find evidence of crosstalk between heterochromatin integrity and RNP granule formation, suggesting the involvement of common limiting factors in control of both transcripts released from polysomes during stress, and those accumulating as a result of de-repression of heterochromatin. Consistent with this, RNA-FISH analyses revealed that, when heterochromatin is disrupted, absence of Cpn1 is associated with hyperaccumulation of non-coding pericentromeric transcripts that localise *in cis* at centromeres. Together, our findings unveil a role for Cpn1 in supporting efficient heterochromatin establishment by enabling the removal of excess heterochromatic transcripts.

## Results

### Efficiency of heterochromatin establishment is influenced by parental genetic background

To investigate factors influencing heterochromatin establishment, we conducted crosses with different CLRC and/or RNAi mutant strains deficient in heterochromatin, and examined the efficiency of heterochromatin re-establishment in the wild-type progeny. An *ade6^+^* reporter gene inserted into the heterochromatic outer repeats of centromere one (*cen1:ade6^+^*) was employed as a readout: in cells lacking centromeric heterochromatin this reporter is expressed resulting in pale/white colonies, whereas assembly of heterochromatin causes silencing of the reporter resulting in red colonies (Figure 1A). When a reporter-carrying strain lacking heterochromatin (e.g. *rik1Δ ago1Δ* double mutant, defective for both H3K9 methylation and RNAi), was crossed to a wild-type strain without the reporter, the resulting wild-type (*rik1+ ago1+*), reporter-carrying progeny were 100% red, consistent with the expected re-establishment of heterochromatin and silencing in this background (Figure 1B). Interestingly, however, when we crossed together two different heterochromatin-deficient strains, e.g. *rik1Δ* and *ago1Δ* single mutants, we observed a mix of red and white colonies in the wild-type progeny (∼80% red, 20% white). Similar results were seen when crossing other pairs of heterochromatin-deficient strains, although the ratios of red:white colonies varied with parental genotype (Figure S1A). In contrast, the ratio of red:white colonies was highly consistent amongst crosses of different, independent *ago1Δ* and *rik1Δ* strains (Figure S1B). It was also consistent regardless of the parent of origin of the reporter (Figure 1B). Passaging of progeny colonies revealed that while the red, silenced state is stable, the white phenotype tends to switch to red over time, consistent with slow heterochromatin re-establishment (Figure 1C). Together these results indicate that, upon reintroduction of missing heterochromatin-related factors by crossing, the efficiency of heterochromatin establishment is dependent on the specific parental background.

**Figure 1.**
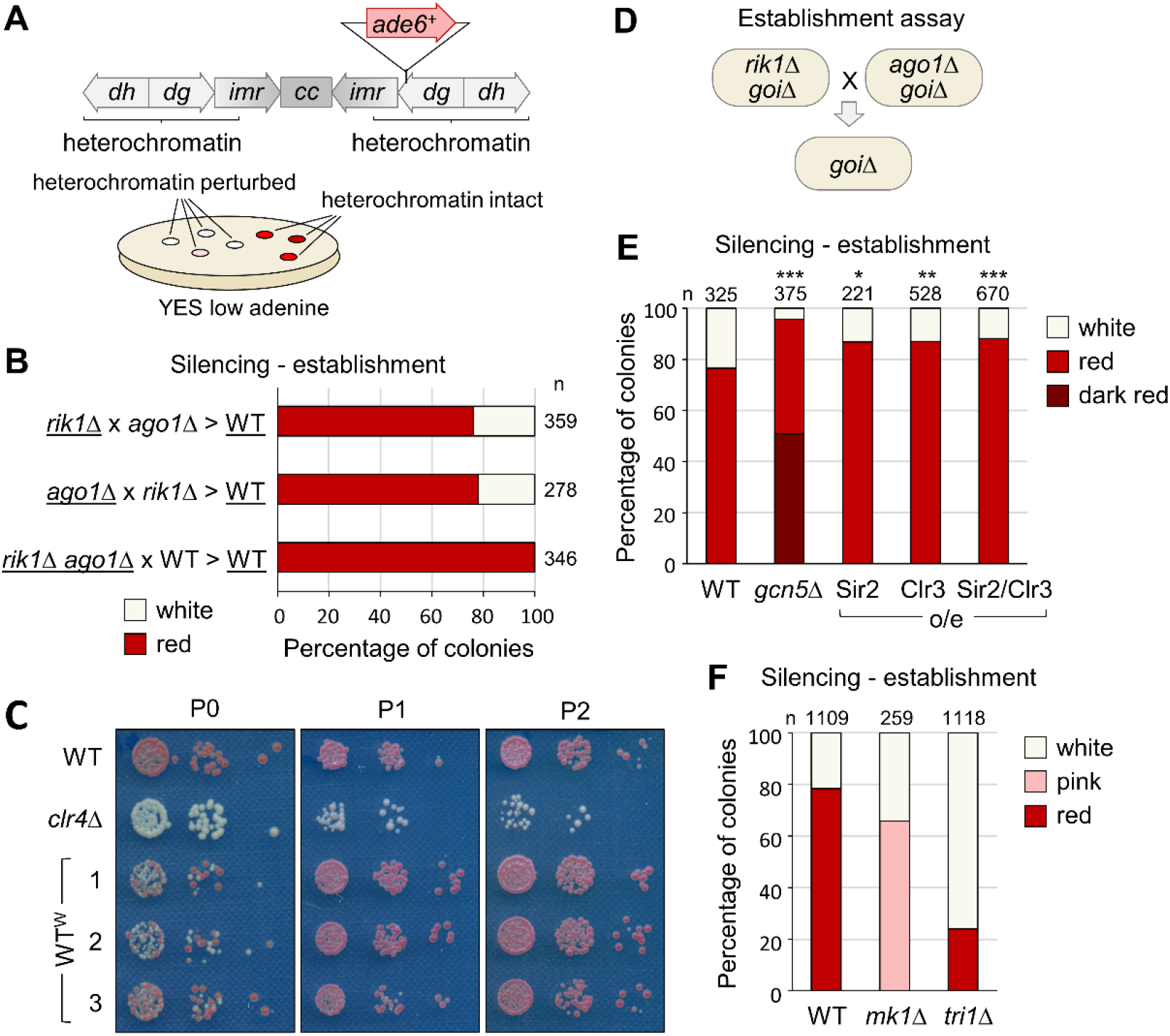
Efficiency of heterochromatin re-establishment is influenced by genetic background. (A) Schematic representation of the *cen1:ade6^+^* reporter system. The position of the *ade6^+^* insertion at centromere one is shown relative to the centromeric outer repeats (*dg* and *dh*), innermost repeats (*imr*) and central core (*cc*). Intact heterochromatin is associated with silencing of *cen1:ade6^+^* reporter resulting in red colonies on low adenine media; loss of heterochromatin alleviates silencing giving pink/white colonies. (B) Proportions of red (*cen1:ade6^+^* silenced) versus white (*cen1:ade6^+^* expressed) colonies in the wild-type progeny of crosses between parental strains of the indicated genotypes (underlining denotes strains carrying the *cen1:ade6^+^* reporter), based on analysis of *n* colonies. (C) Passaging of wild-type ‘white’ (WT^W^) colonies selected from B. (D) Schematic of the cross-based establishment assay. (E) and (F) Proportions of red (*cen1:ade6^+^* silenced) versus pink/white (*cen1:ade6^+^* expressed) colonies in the otherwise wild-type progeny of *rik1Δ* x *ago1Δ* crosses performed in the indicated genetic backgrounds, based on analysis of *n* colonies. Relative to wild-type, asterisks denote p ≤ 0.05 (*), p ≤ 0.01 (**) or p ≤ 0.001 (***) from chi-squared (χ^2^) test analysis.

We next tested whether the rate of heterochromatin re-establishment in the progeny resulting from a cross between *ago1Δ* and *rik1Δ* strains could be modulated by backgrounds expected to promote or suppress heterochromatin assembly. Since histone deacetylation is required to enable stable H3K9 methylation, we reasoned that the rate of heterochromatin establishment might be increased by interventions that reduce histone H3 acetylation. To test this, we repeated the cross in backgrounds either lacking the histone acetyltransferase Gcn5 (36,37), or over-expressing deacetylases Sir2 and/or Clr3. For example, we deleted *gcn5^+^* in *ago1Δ* and *rik1Δ* strains, crossed the resulting strains together, and analysed the *gcn5Δ ago1+ rik1+* progeny (Figure 1D). As predicted, we found that removal of Gcn5 was sufficient to increase the efficiency of heterochromatin establishment, resulting in the proportion of red colonies in the cross progeny increasing to almost 100% (Figure 1E). Over-expression of Sir2 and/or Clr3 also resulted in a smaller but still significant increase in red colony proportion, consistent with their reported roles in promoting heterochromatin establishment (32,33). Conversely, we also tested the impact of deleting two factors previously shown to promote heterochromatin establishment, Mkt1 and Tri1 (34,35). As expected, in the absence of either of these factors the efficiency of heterochromatin establishment appeared reduced, with *tri1Δ* progeny showing a strong reduction in the proportion of red colonies, and *mkt1Δ* progeny showing a mixture of white and pink but no red colonies (Figure 1F). Thus, this cross-based system can serve as a sensitized assay to detect and quantify changes in efficiency of heterochromatin establishment.

### Cpn1 is a novel factor required for efficient heterochromatin establishment

As a starting point to uncover potential novel heterochromatin establishment factors, we returned to data from a previous genetic screen for factors influencing heterochromatin maintenance (38). In that screen, maintenance of silencing of the *cen1:ade6^+^* reporter was primarily assessed via colony growth on media lacking adenine, which was scored semi-quantitatively on a scale of one to four, where four indicated expression comparable to wild-type, and one indicated strong de-repression. 75 mutant strains were given a growth score of three, suggesting possible mild de-repression of the reporter, but falling below the threshold (of two) set for further follow-up analysis as a candidate maintenance factor. We reasoned that these strains showing potential slight defects in heterochromatin maintenance might include deletions of factors that play a greater role in heterochromatin establishment. To further narrow down the list we selected factors with no known *S. cerevisiae* ortholog (according to PomBase (39)), since H3K9 methylation and RNAi are absent in this species. This resulted in a shortlist of 13 genes, one of which was known establishment factor Tri1, validating the approach. Of the remaining 12 genes, three were previously uncharacterised, and of these one in particular stood out: SPAC12G12.07c, hereon named *cpn1^+^* for reasons outlined below. Like another known establishment factor, Mkt1, the product of this gene has previously been reported to physically associate with components of the MTREC exosome adaptor complex that is implicated in maintenance of facultative heterochromatin domains (40), making it a strong candidate as a novel pathway component. To test this, we assessed the impact of *cpn1^+^* deletion in our *rik1Δ* x *ago1Δ* establishment assay. Strikingly, *cpn1Δ* progeny showed a clear reduction in the proportion of red colonies, similar to what was seen in the *tri1Δ* background, suggesting that the product of this gene is required for efficient heterochromatin establishment (Figure 2A).

**Figure 2.**
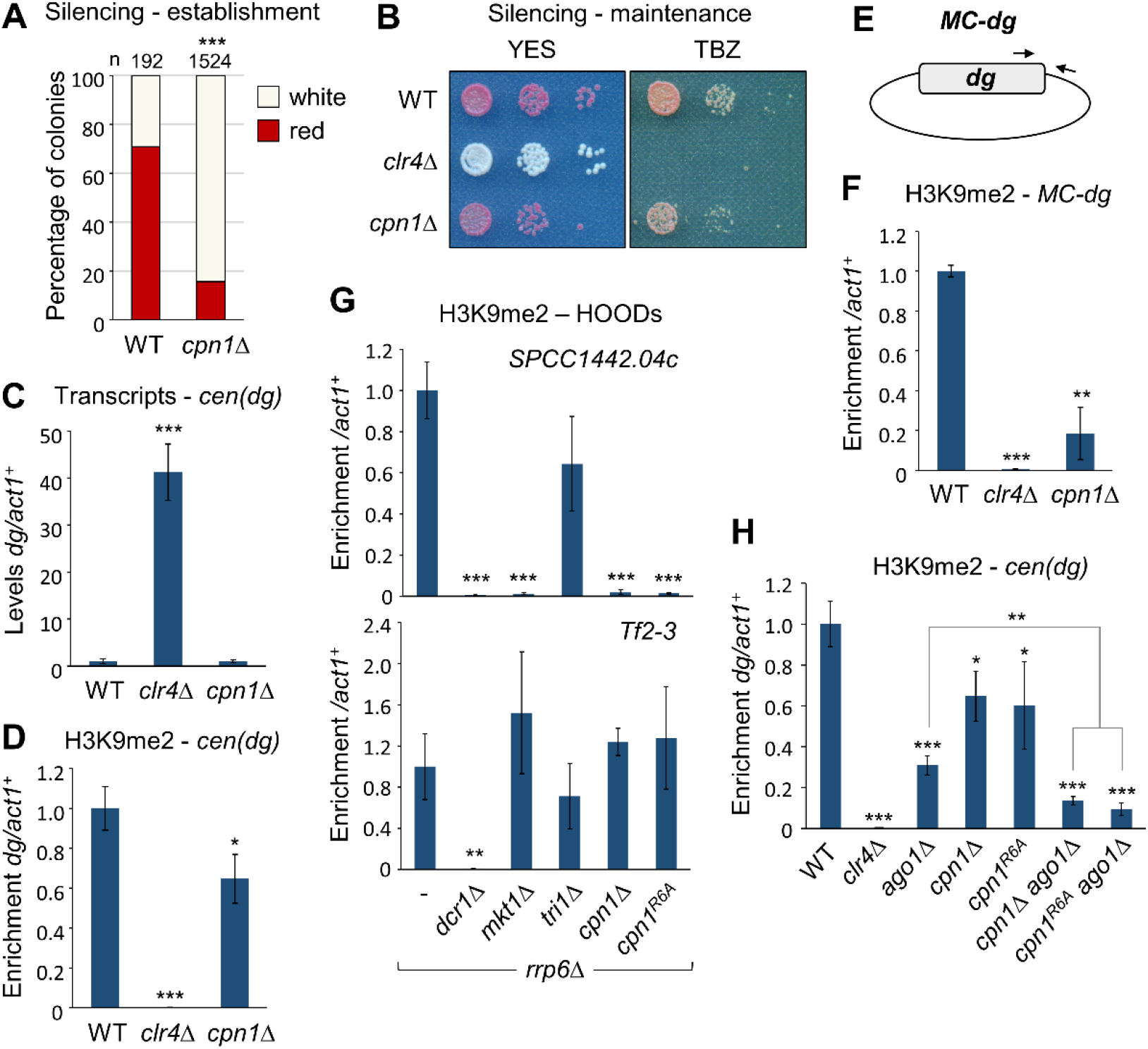
Cpn1 is a novel factor required for efficient heterochromatin establishment. (A) Assay for establishment of silencing of the *cen1:ade6^+^* reporter. Shown are proportions of red (*ade6^+^* silenced) versus white (*ade6^+^* expressed) colonies in the otherwise wild-type progeny of *rik1Δ* x *ago1Δ* cross performed in wild-type or *cpn1Δ* background (as in 1D), based on analysis of *n* colonies. Asterisks denote p ≤ 0.001 (***) from chi-squared (χ^2^) test analysis. (B) Assay for maintenance of silencing of the *cen1:ade6^+^* reporter (red colonies indicate silencing; white colonies loss of silencing), and for sensitivity to TBZ (20 μg/μl). (C) RT-qPCR analysis of *cen*(*dg*) transcript levels relative to *act1^+^*, normalised to wild-type. (D) ChIP-qPCR analysis of H3K9me2 levels at *cen*(*dg*) relative to *act1^+^*, normalised to wild-type. (E) Schematic of the minichromosome (*MC-dg*) plasmid. (F) ChIP-qPCR analysis of H3K9me2 levels established *de novo* on the *MC-dg* plasmid, relative to *act1^+^*, normalised to wild-type. (G) ChIP-qPCR analysis of H3K9me2 levels at two previously described HOODs, *SPCC1442.04c* and *Tf2-3,* relative to *act1^+^*, normalised to wild-type. (H) ChIP-qPCR analysis of H3K9me2 levels at *cen*(*dg*) relative to *act1^+^*, normalised to wild-type. In each case data are averages of three biological replicates and error bars represent one SD. Asterisks denote p ≤ 0.05 (*), p ≤ 0.01 (**) or p ≤ 0.001 (***) from Student’s t-test analysis.

Our previous screen results had suggested that deletion of *cpn1^+^* might result in a small defect in heterochromatin maintenance (38). To assess this in more detail, we deleted *cpn1^+^* in a wild-type strain carrying the *cen1:ade6^+^* reporter to assess the effect on heterochromatin maintenance. Colony colour of *cpn1Δ* strains was indistinguishable from wild-type, suggesting little or no effect on maintenance of reporter gene silencing (Figure 2B). Consistent with this, *cpn1Δ* strains, in contrast to *clr4Δ* strains, showed little increase in sensitivity to the microtubule destabilising drug TBZ, indicating that centromere function is maintained. This was further supported by RT-qPCR analysis that showed no increase in accumulation of non-coding pericentromeric (*dg*) transcripts (Figure 2C). In addition, ChIP-qPCR analysis of H3K9me2 revealed only a small reduction in H3K9 methylation levels at pericentromeric repeat sequences (Figure 2D). Hence loss of Cpn1 function has little effect on heterochromatin maintenance, and rather appears to particularly affect heterochromatin establishment.

As a complementary approach to further verify and characterise the heterochromatin establishment defects in *cpn1Δ* cells, we employed an alternative assay whereby we assessed *de novo* heterochromatin establishment on a newly introduced minichromosome, as described previously (32,35). Wild-type and mutant strains were transformed with a minichromosome plasmid (*MC-dg*) that carries a 5.6kb portion of centromeric (*dg*) outer-repeat sequence that is targeted by endogenous centromeric siRNAs (Figure 2E). ChIP-qPCR analysis revealed that, whereas in wild-type cells, heterochromatin is efficiently established on the plasmid resulting in high levels of H3K9me2, in cells lacking Cpn1 the levels of H3K9me2 established on the plasmid are substantially reduced (Figure 2F). This is consistent with the reduced efficiency of re-initiation of pericentromeric reporter gene silencing observed in *cpn1Δ* cells, and supports the conclusion that absence of Cpn1 is associated with defective heterochromatin establishment.

Cpn1 has previously been reported to interact with MTREC components including Red1, Red5, Mtl1 and Pir2 (40). We therefore tested whether Cpn1 also functions together with MTREC in facilitating assembly of facultative heterochromatin. Two classes of MTREC-dependent facultative heterochromatin domains can be distinguished: heterochromatin islands, which are RNAi-independent and typically serve to silence meiotic genes during vegetative growth (17,18), and so-called HOODs, which are dependent on both MTREC and RNAi and form at loci such as transposons in the absence of Rrp6 (19). In the case of heterochromatin islands, RT-qPCR analysis revealed that, while absence of MTREC component Red1 results in loss of silencing of meiotic genes such as *mei4^+^* and *ssm4^+^*, the removal of Cpn1 has no effect at these loci (Figure S2A). Similarly, at HOODs (assessed in *rrp6Δ* background), ChIP-qPCR analysis indicated that, whereas H3K9 methylation is lost upon deletion of *dcr1^+^*, it is retained at most loci tested, including Tf2-3, in the absence of *cpn1^+^* (Figure 2G and S2B). However, the one exception to this is the HOOD associated with meiotic gene SPCC1442.04c, where we found heterochromatin to be strongly dependent on functional Cpn1 (Figure 2G). This was striking since we previously found that this particular HOOD (but not other HOODs or islands) is also dependent on Mkt1 (but not Tri1) (35). This HOOD is also unusual in being Red1-independent (19). Hence Cpn1, like Mkt1, plays a role in supporting RNAi-dependent facultative heterochromatin formation in certain contexts, possibly acting in parallel to Red1.

The commonalities described above prompted us to probe whether Mkt1 and Cpn1 might be functionally linked. We showed previously that Mkt1 acts in the same pathway as RNAi for pericentromeric silencing. To test if this is also true for Cpn1, we assessed pericentromeric H3K9me2 levels in *cpn1Δ ago1Δ* double mutant cells. In contrast to our previous observations for Mkt1, we found that the *cpn1Δ ago1Δ* double mutants displayed additive defects in H3K9 methylation as compared to either single mutant, suggesting independent effects (Figure 2H). Hence it appears that while Mkt1 functions together with RNAi, Cpn1 functions in another, separate pathway.

### Cpn1 is a CAPRIN family protein

At the time of this study no orthologs of the SPAC12G12.07c gene product had been identified outside of fungi. However, through an iterative search for remote homology using JACKHMMER (41), we identified 1:1 orthology with human protein CAPRIN1. Although they have only 15% identity at the amino level, the human and fission yeast proteins share similar motif organisation, including N-terminal coiled-coil and C-terminal RG/RGG rich regions (Figure 3A). Moreover, comparison of structural models generated by AlphaFold (42,43) also reveals clear similarities in structural organisation (Figure 3B). We therefore named the SPAC12G12.07c gene product as fission yeast Caprin protein, Cpn1.

**Figure 3.**
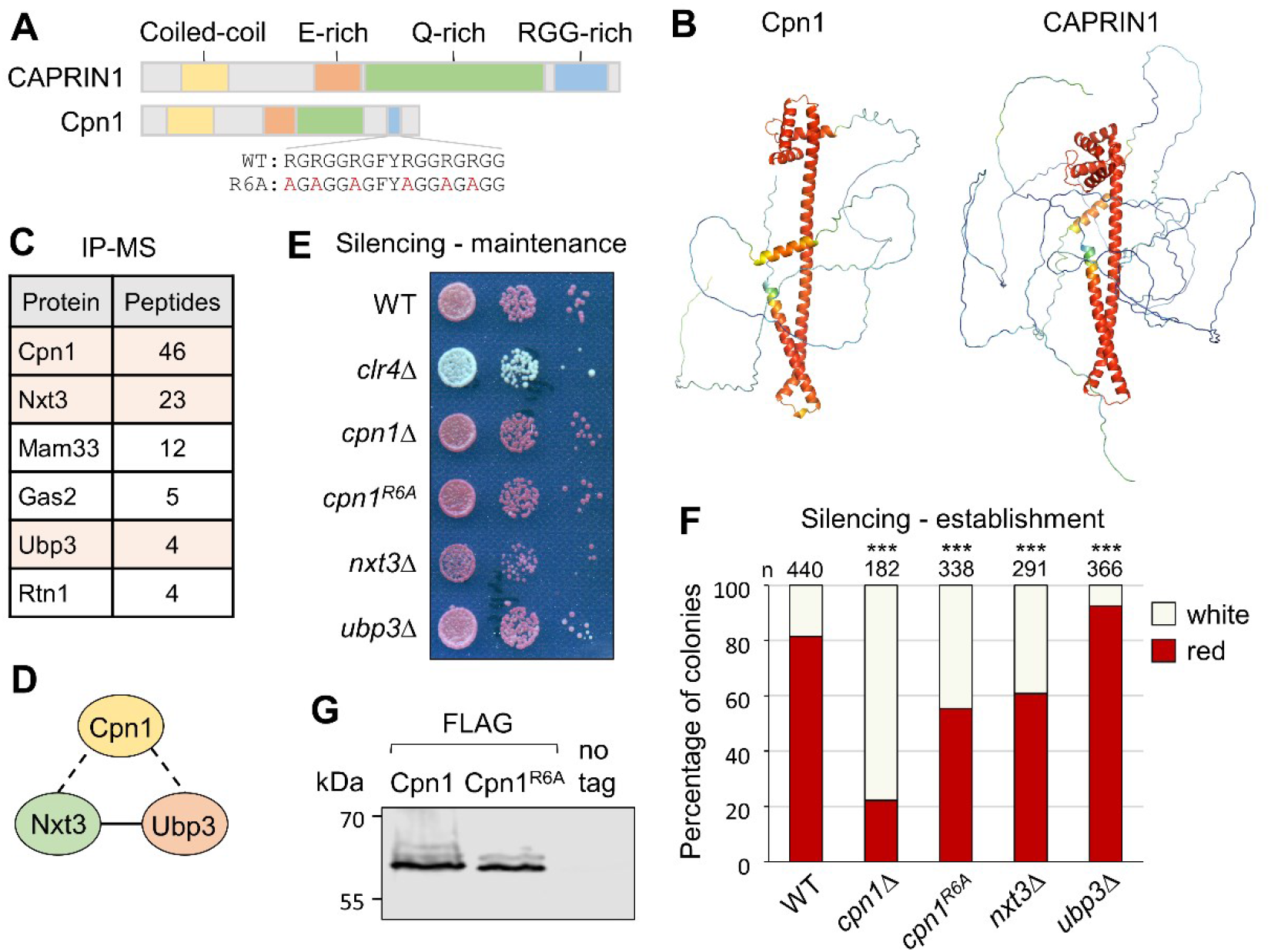
Cpn1 is a CAPRIN family protein. (A) Schematic representation of CAPRIN proteins in human (CAPRIN1) and *S. pombe* (Cpn1). (B) Alphafold structural models of Cpn1 and CAPRIN1. (C) List of proteins found specifically and reproducibly associated with Cpn1-FLAG by affinity purification and mass spectrometry (LC-MS/MS). Shown are total peptide counts for all proteins represented by ≥ 2 peptides in purifications from Cpn1-FLAG strains, but not untagged control strains, in two independent experiments. (D) Putative Cpn1 complex (dotted lines denote interactions detected here; solid line indicates interaction detected previously by IP-western (45)). (E) Assay for maintenance of silencing of *cen1:ade6^+^* reporter: red colonies on low adenine media indicate silencing, and white colonies absence of silencing. (F) Assay for establishment of silencing of *cen1:ade6^+^* reporter: shown are proportions of red (*ade6^+^* silenced) versus white (*ade6^+^* expressed) colonies in the otherwise wild-type progeny of *rik1Δ* x *ago1Δ* crosses performed in the indicated deletion backgrounds (as in 1D), based on analysis of *n* colonies. Relative to wild-type, asterisks denote p ≤ 0.001 (***) from chi-squared (χ^2^) test analysis. (G) Western blot analysis of affinity-purified FLAG-tagged wild-type Cpn1 and Cpn1^R6A^.

Human CAPRIN1 is a ubiquitously expressed mRNA-binding protein that is known to be involved in regulation of stress granule formation in association with two partner proteins, G3BP1 and USP10 (44). To investigate fission yeast Cpn1 interaction partners, we C-terminally FLAG-tagged Cpn1 at the endogenous locus and performed immunoprecipitation followed by liquid chromatography-tandem mass spectrometry (LC-MS/MS) to identify associated proteins. Only five proteins were specifically and reproducibly identified in precipitates from Cpn1-FLAG-expressing cells but not controls, and strikingly, two of these were Nxt3 and Ubp3, the fission yeast orthologs of human CAPRIN1 partner proteins G3BP1 and USP10, respectively (Figure 3C and D). Nxt3 and Ubp3 are themselves known to interact, and to localise to stress granules (45,46). The association with Nxt3 and Ubp3 further supports the conclusion that fission yeast Cpn1 is the functional ortholog of human CAPRIN1.

To explore whether Nxt3 and Ubp3 also function together with Cpn1 to influence heterochromatin assembly, we generated *nxt3Δ* and *ubp3Δ* deletion strains and tested them in heterochromatin maintenance and establishment assays. While absence of these partner proteins, like loss of Cpn1 itself, had little effect on heterochromatin maintenance (Figure 3E), it did affect heterochromatin establishment in our *rik1Δ* x *ago1Δ*. Interestingly, the two deletions had opposing effects: while in *nxt3Δ* cells, as in *cpn1Δ* cells, the efficiency of heterochromatin establishment was reduced (lower proportion of red colonies in the cross progeny compared to wild-type), in *ubp3Δ* cells we conversely saw a small increase in establishment efficiency (increased proportion of red colonies; Figure 3F). Considering what is known about their human orthologs, this may be explained by opposing functions of these proteins, since CAPRIN1 and G3BP1 are both RNA-binding proteins that have been shown act cooperatively to promote stress granule assembly, while USP10, which lacks an RNA-binding domain, acts to limit assembly (44). Hence Cpn1 acts together with its interaction partners to influence heterochromatin establishment efficiency.

The RGG-rich region in human CAPRIN1 has been shown to be important for RNA-binding and stress granule formation (47). To assess whether the RG/RGG-rich motif in fission yeast Cpn1 is also required for its function in heterochromatin establishment, we generated strains in which each of the six arginine residues in this region were mutated to alanine (Cpn1^R6A^; Figure 3A). Testing this mutant in our *rik1Δ* x *ago1Δ* establishment assay, we observed a reduction in the proportion of red colonies in the progeny compared to wild-type, indicating reduced efficiency of heterochromatin establishment, although the effect was not as strong as that associated with *cpn1^+^* deletion (Figure 3F). Moreover, in Cpn1^R6A^-expressing cells we also observed loss of H3K9me2 from the SPCC1442.04c HOOD locus, and a reduction in H3K9me2 maintenance at pericentromeric repeats in *ago1Δ* background, comparable to that in *cpn1Δ* cells (Figure 2G and H). Western blot analysis of FLAG-tagged wild-type and mutant Cpn1 proteins showed no difference in protein accumulation (Figure 3G), confirming that the mutations do not affect protein stability. Hence the RGG-rich region contributes to Cpn1 function in facilitating heterochromatin assembly.

Since the RGG motifs in human CAPRIN1 are involved in RNA binding, and since we have shown previously that Mkt1 interacts with non-coding pericentromeric transcripts (35), we tested whether Cpn1 might also bind to pericentromeric RNAs. As previously, we performed RNA-IPs in cells lacking Sir2, to relieve transcriptional repression at pericentromeric heterochromatin (35). We detected specific enrichment of pericentromeric transcripts (*dg* and *imr*) in Mkt1-FLAG pulldowns relative to controls, consistent with our previous results. However, no such enrichment was observed in Cpn1-FLAG pull-downs, suggesting that Cpn1 does not stably bind these RNAs (Figure S3). We also tested whether association of Mkt1 with pericentromeric transcripts is dependent on Cpn1, but again found no evidence of this, with both *dg* and *imr* transcripts being similarly enriched in Mkt1-FLAG pull downs in the presence and absence Cpn1 (Figure S3). This supports the conclusion that Cpn1 and Mkt1 function independently in heterochromatic silencing.

### Cpn1 promotes stress granule formation

To test whether Cpn1, like its human ortholog, localises to stress granules, we introduced a C-terminal GFP tag at the endogenous locus and performed live cell imaging analyses. While in unstressed cells we saw a diffuse pattern of localisation, in cells subjected to stress (heat shock or glucose starvation), Cpn1 could be seen to accumulate in discrete cytoplasmic foci. Analysis of cells co-expressing stress granule marker poly(A)-binding protein Pabp tagged with RFP (48) indicated that Cpn1 colocalises with Pabp, confirming that these foci correspond to stress granules (Figure 4A). Similar results were obtained for GFP-tagged Cpn1^R6A^, indicating that the RGG-rich region is not required for Cpn1 localisation to stress granules. We also generated strains expressing C-terminally GFP-tagged Nxt3 or Ubp3, and confirmed that both of these proteins also colocalise with both Pabp and Cpn1 in stress-induced granules (Figure 4B and C). Western blot and heterochromatin establishment assays confirmed that all of the GFP-tagged proteins are stably expressed and functional (Figure S4). Hence Cpn1, together with its partner proteins Nxt3 and Ubp3, does indeed partition into phase-separated, membrane-less organelles in response to stress.

**Figure 4.**
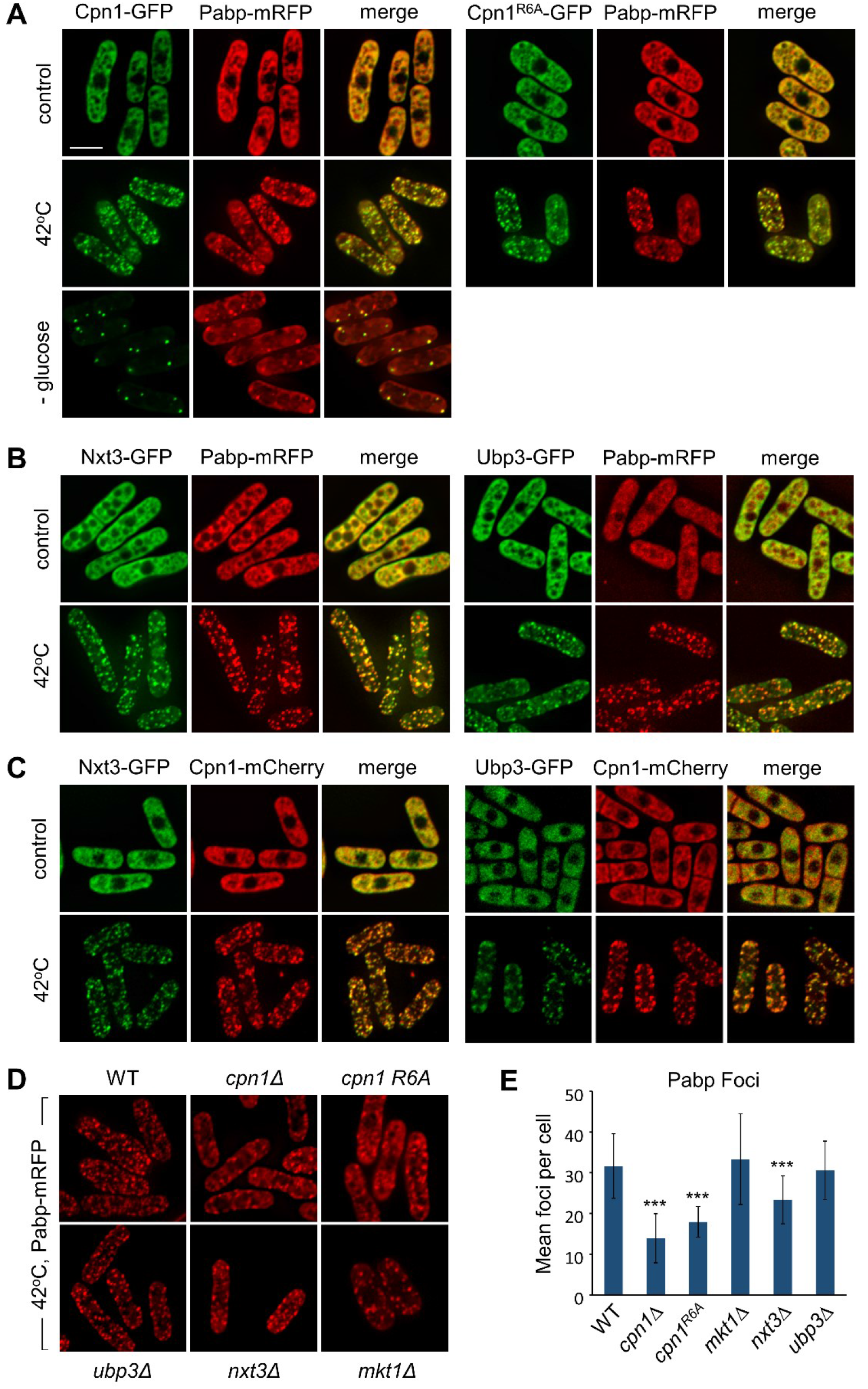
Cpn1 localises to stress granules and promotes their formation. (A) Representative images from two-colour live-cell imaging of Cpn1-GFP or Cpn1^R6A^-GFP (green), and Pabp-mRFP (red), in cells grown at 32°C (control) and exposed to 42°C heat shock for 20 min, or glucose starvation for 20 min. Bar indicates 6 µm. (B and C) Representative images from two-colour live-cell imaging of Nxt3-GFP or Ubp3-GFP (green), and either Pabp-mRFP (red; B) or Cpn1-mCherry (red; C), in cells either untreated or treated with heat shock as above. (D) Representative images from live-cell imaging of Pabp-mRFP (red) in the indicated genetic backgrounds in cells exposed to heat shock as above. (E) Quantification of the mean number of Pabp-mRFP foci per cell from the imaging analysis shown in D (*n* = 100). Error bars represent one SD and asterisks denote p ≤ 0.001 (***) from Student’s t-test analysis.

Human CAPRIN1 has also been shown to be required to promote stress granule assembly, in a manner dependent on its RGG repeats (44,47). To test if the same is true for fission yeast Cpn1, we quantified Pabp focus formation in response to stress in cells lacking Cpn1 or other factors. In line with predictions, in cells lacking Cpn1 we found that the average number of Pabp foci formed per cell following heat shock was substantially reduced (Figure 4D and E). Granule formation was also reduced in cells expressing Cpn1^R6A^, suggesting that the RGG-rich motif is important for Cpn1 function in stress granule assembly. We also observed a reduction in granules in cells lacking Nxt3, consistent with findings for the human ortholog G3BP1 (44). In contrast, no change in Pabp focus formation was observed in cells lacking Ubp3, or Mkt1 (Figure 4D and E). Hence like their human orthologs, Cpn1 and Nxt3 are required to promote stress granule formation.

### Crosstalk between heterochromatin integrity and cytoplasmic RNP granule formation

We next set out to explore the possible connection between assembly of heterochromatin and RNP granules. First, we tested whether mutations that disrupt heterochromatin impact on granule formation. Interestingly, whereas in the wild-type background, Pabp foci are not seen in unstressed cells, when we deleted genes encoding components of either the RNAi machinery (Ago1) or the CLRC H3K9 methyltransferase complex (Clr4 or Rik1), we observed a proportion of cells exhibiting Pabp foci even in the absence of stress (Figure 5A and B). This was most prominent in *ago1Δ* cells, where granules were visible in approximately 15% of cells. Analysis of cells co-expressing Cpn1-GFP confirmed that Cpn1 also localises to these granules, as for canonical stress granules (Figure 5C; colocalization was seen in all cells displaying visible granules). We further tested whether the formation of these *ago1Δ* or *clr4Δ*-dependent granules is dependent on Cpn1. Indeed, we found that granule formation was greatly reduced in *cpn1Δ ago1Δ* or *cpn1Δ clr4Δ* double mutants compared to the respective single mutants, indicating that Cpn1 is required to promote the formation of these heterochromatin deficiency-induced granules (Figure 5B).

**Figure 5.**
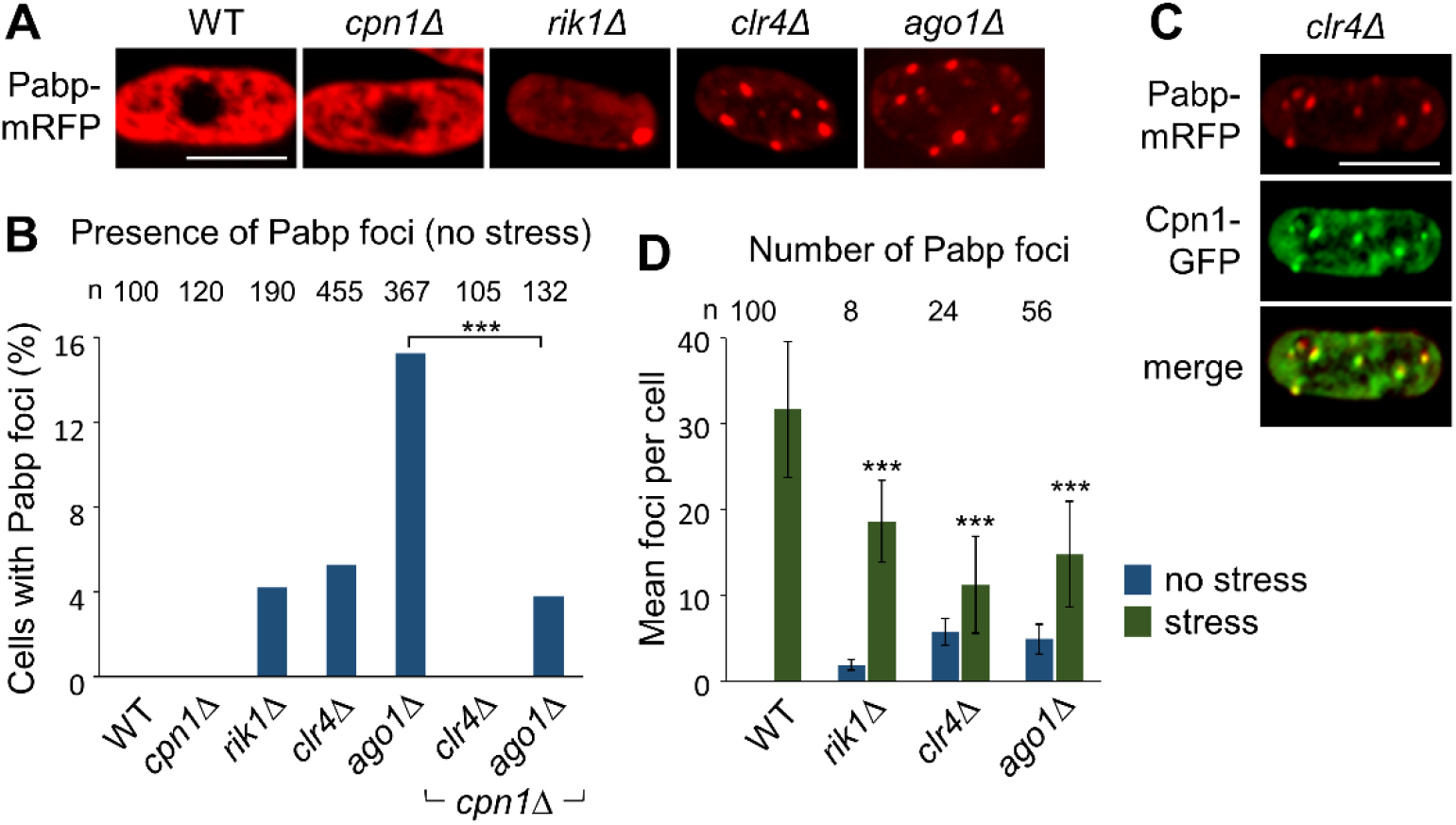
Disruption of heterochromatin alters the formation of Pabp-containing RNP granules. (A) Representative images from live-cell imaging of Pabp-mRFP (red) in the indicated genetic backgrounds in unstressed cells. Bar indicates 6 µm. (B) Quantification of the proportion of unstressed cells displaying Pabp-mRFP foci from the imaging analysis shown in A, based on analysis of *n* cells. Asterisks denote p ≤ 0.001 (***) from chi-squared (χ^2^) test analysis. (C) Representative images from two-colour live-cell imaging of Cpn1-GFP (green), and Pabp-mRFP (red), in unstressed *clr4Δ* cells. (D) Quantification of the mean number of Pabp-mRFP foci per cell in the indicated genetic backgrounds in unstressed cells (based on analysis of *n* cells containing foci), or cells exposed to 42°C heat shock for 20 min (*n* = 100; virtually all cells contained foci). Error bars represent one SD and asterisks denote p ≤ 0.001 (***) from Student’s t-test analysis.

We also tested what happens in heterochromatin deficient cells in the presence of stress. Interestingly, we saw an opposite effect: whereas heterochromatin mutants show increased Pabp granule formation in absence of stress, they show a significant reduction in the average number of Pabp foci formed in the presence of stress, compared to wild-type cells (Figure 5D). This suggests that RNP granules observed in the absence of heterochromatin may be assembled at the expense of those formed during stress, indicating an element of competition that could reflect involvement of one or more common limiting factors.

### Cpn1 helps limit accumulation of heterochromatic transcripts on chromatin

We hypothesised that the RNP granules that are occasionally formed in response to heterochromatin disruption may be linked to sequestration and/or degradation of excess heterochromatic transcripts that accumulate in the absence of heterochromatin-associated silencing. While previous fractionation analyses have provided evidence that heterochromatic transcripts tend to be retained on chromatin (24), to our knowledge there has been no direct visualisation of heterochromatic transcript subcellular localisation in *S. pombe*. To explore whether the RNP granules formed in response to heterochromatin disruption might harbour non-coding pericentromeric RNAs, we therefore set out determine the subcellular localisation of these RNAs by single-molecule mRNA fluorescence *in situ* hybridization (smRNA-FISH) using probes targeting *cen*(*dg*) transcripts. Previous RT-qPCR and northern blot analyses have indicated that pericentromeric transcript levels are low in wild-type cells, but greatly elevated in heterochromatin mutants where transcriptional silencing is alleviated (6,10,49,50). Consistent with this, by smRNA-FISH we could detect a strong signal for *cen*(*dg*) transcripts in *clr4Δ* cells, but not in wild-type cells, confirming that the signal is *dg* transcript-specific (Figure 6A). Notably, *cen*(*dg*) transcripts were detected in one strong focus in the nucleus, which we suspected might corresponded to the position of centromeres, which tend to cluster together at the nuclear periphery in *S. pombe* (51). To test this, we combined smRNA-FISH with immunofluorescence analysis of GFP-tagged centromeric histone variant CENP-A^Cnp1^. The *cen*(*dg*) transcript focus was consistently found adjacent to, or overlapping, the CENP-A ^Cnp1^ focus, consistent with transcripts being primarily retained at the pericentromeres (Figure 6B).

**Fig 6:**
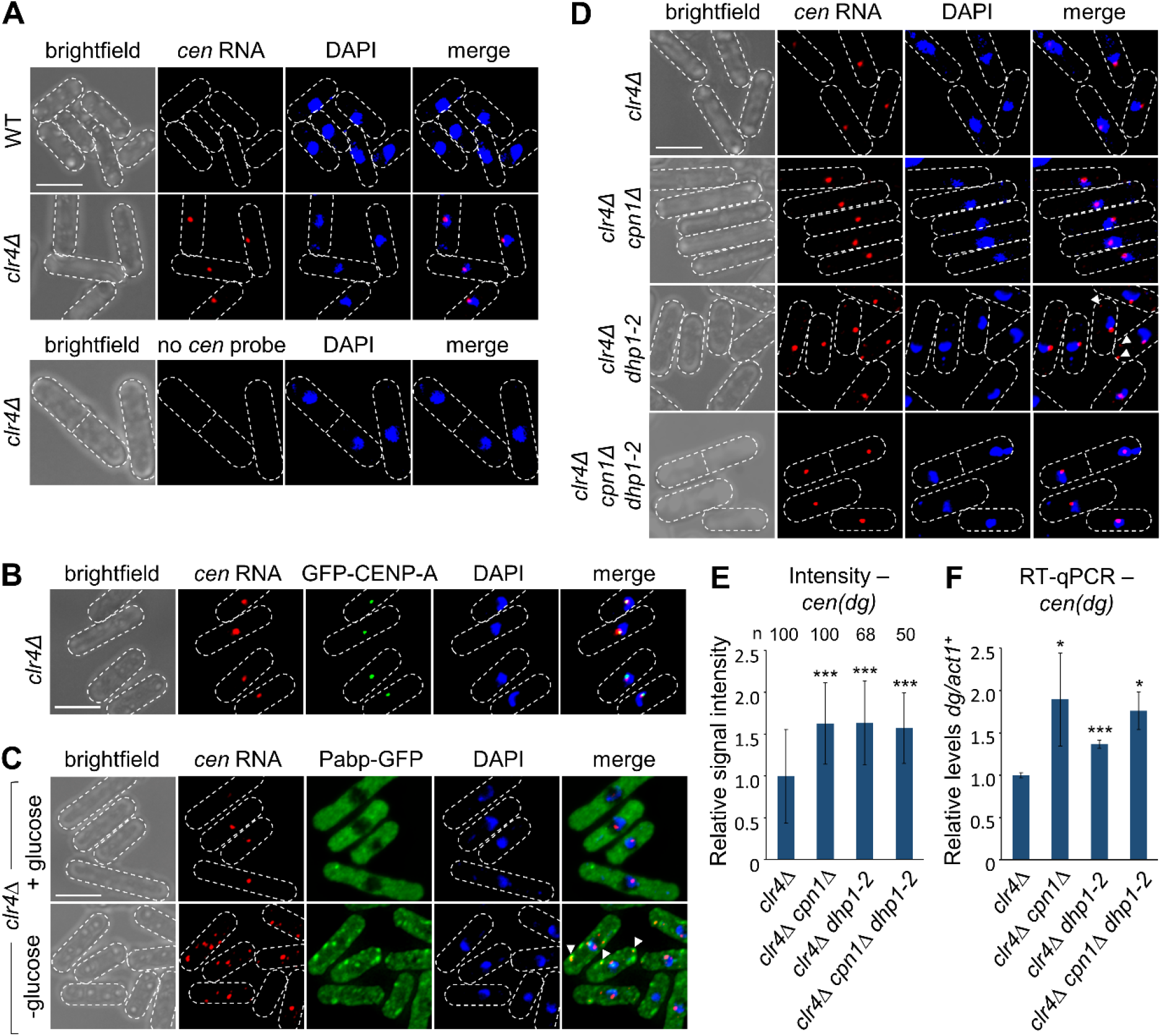
Absence of Cpn1 results in increased accumulation of heterochromatic transcripts at centromeres in heterochromatin deficient cells. (A) Representative images from smRNA-FISH analysis of pericentromeric *(cen(dg))* RNA in wild-type or *clr4Δ* strains. Bar indicates 6 µm. (B) Representative images from simultaneous analysis of *cen(dg)* RNA by smRNA-FISH, and GFP-CENP-A^Cnp1^, in *clr4Δ* cells. (C) Representative images from simultaneous analysis of *cen(dg)* RNA by smRNA-FISH, and Pabp-GFP, in *clr4Δ* cells either unstressed (+ glucose) or stressed by 20 min of glucose starvation (-glucose). Arrow heads highlight examples of cytoplasmic co-localisation (Pearson’s correlation coefficient: 0.42). (D) Representative images from smRNA-FISH analysis of *cen(dg)* RNA in *clr4Δ*, *clr4Δ cpn1Δ*, *clr4Δ dhp1-2*, and *clr4Δ cpn1Δ dhp1-2* cells. Arrow heads highlight examples of cytoplasmic RNA foci. (E) Quantification of the mean signal intensity for nuclear *cen(dg)* RNA foci, normalised to *clr4Δ* cells, from the smRNA-FISH analysis shown in D. (F) RT-qPCR analysis of total cellular levels of *cen(dg)* RNA relative to *act1^+^*, normalised to *clr4Δ* cells. RT-qPCR data are averages of three biological replicates. In all cases, error bars represent one SD, and asterisks denote p ≤ 0.05 (*), or p ≤ 0.001 (***), from Student’s t-test analysis.

To address whether *cen*(*dg*) transcripts might also localise to the Pabp foci seen in a small proportion of (unstressed) heterochromatin-deficient cells, we combined RNA-FISH with immunofluorescence analysis of Pabp-GFP. However, in either *clr4Δ* or *ago1Δ* cells, where visible Pabp foci were formed we could not detect co-localisation of *cen*(*dg*) transcripts, suggesting that these foci are not sites of stable heterochromatic transcript accumulation (Figure 6C and S5A, upper panels). It remained possible that these foci might rather be associated with RNA degradation, so to further explore the potential for heterochromatic transcripts to be translocated from the nucleus to sites of Cpn1 aggregation in the cytoplasm, we performed additional RNA-FISH analyses in *clr4Δ* or *ago1Δ* cells subject to stress, to induce formation of stress granules. Notably, following glucose starvation, additional *cen*(*dg*) transcript foci were observed, at least some of which appeared to colocalise with cytoplasmic stress granules (Figure 6C and S5A, lower panels). It may be that this cytoplasmic localisation is only detectable in stress as this causes redistribution of *cen*(*dg*) RNAs, via their associated RBPs, into more stable condensates. Hence heterochromatic transcripts can exit the nucleus, and co-localise with Cpn1 and associated factors in the cytoplasm.

To assess whether Cpn1 impacts pericentromeric transcript localisation and/or accumulation in the absence of stress, we performed further RNA-FISH analyses in cells lacking Cpn1. Interestingly, we observed a visible increase in the intensity of the *cen(dg)* transcript signal at centromeres in *clr4Δ cpn1Δ* double mutants compared to *clr4Δ* single mutants, suggesting that Cpn1 is required to help limit accumulation of these transcripts at centromeres (Figure 6D and E). The same effect was also seen in *ago1Δ* background (Figure S5B and C). In principle, this Cpn1-dependent protection of centromeres from excess RNA accumulation might be achieved either via sequestration of these RNAs elsewhere in the cell, or through their degradation. To distinguish between these two possibilities, we analysed total cellular levels of *cen(dg)* transcripts by RT-qPCR. Strikingly, this revealed that deletion of *cpn1^+^* in either *clr4Δ* or *ago1Δ* background results in a significant increase in total *cen(dg)* transcripts levels, beyond the already high levels present in absence of Clr4 or Ago1 alone (Figure 6F and S5D). Thus, Cpn1 appears to be required to facilitate degradation of excess heterochromatic transcripts, limiting their accumulation on chromatin.

Finally, we reasoned that if Cpn1 facilitates RNA degradation in the cytoplasm, suppression of degradation (independently of other Cpn1 functions) might lead to visible cytoplasmic accumulation of *cen(dg)* transcripts. Since Caprin proteins do not themselves appear to have nuclease activity, it is likely that Cpn1 primarily influences transcript localisation and/or accessibility for degradation by one or more ribonucleases. We identified the 5’-3’ exoribonuclease Dhp1 (ortholog of human Xrn2) as a likely candidate, since: (1) in human embryonic stem cells, Xrn2 has been implicated in mediating CAPRIN1-dependent degradation of selected transcripts in the cytoplasm (52); and (2) Dhp1, like Cpn1, has been shown to be required for efficient heterochromatin establishment in *S. pombe* (22,23). To explore the influence of Dhp1 on pericentromeric transcript accumulation, we performed further RNA-FISH analyses in *dhp1* mutant cells. Because it is an essential gene, we used a previously described temperature-sensitive allele, *dhp1-2*, that causes defects in heterochromatin, but little impairment of growth, at the permissive temperature (23). Strikingly, whereas in *clr4Δ* cells a single *cen(dg)* transcript focus can be seen in the nucleus, in *dhp1-2 clr4Δ* cells, we observed the appearance of additional *cen(dg)* RNA foci in the cytoplasm (Figure 6D; cytoplasmic RNA foci were detected in ∼22% of *dhp1-2 clr4Δ* cells, possibly reflecting the fact that *dhp1-2* is a hypomorphic allele). Hence, active degradation prevents accumulation of *cen(dg)* transcripts in the cytoplasm in *clr4Δ* cells in normal conditions. Importantly, the cytoplasmic *cen(dg)* RNA foci were abolished in *cpn1Δ dhp1-2 clr4Δ* triple mutant cells, consistent with loss of Cpn1 causing a defect upstream of Dhp1, likely in the localisation of transcripts for degradation. Together, our data suggest that Cpn1 helps to enable heterochromatic transcript degradation, limiting excess accumulation of RNA on chromatin that may otherwise interfere with heterochromatin assembly.

## Discussion

Multiple RNA processing pathways have been found to contribute to RNA-directed heterochromatin assembly and silencing of associated transcripts. Here we have identified a novel role for the fission yeast ortholog of human CAPRIN1, an RNA-binding protein, in facilitating efficient heterochromatin establishment. Absence of Cpn1 has little effect on the maintenance of pre-existing heterochromatin. However, in cells where heterochromatin is abolished and hence non-coding heterochromatic transcripts accumulate, loss of Cpn1 is associated with build-up of even higher levels heterochromatic RNAs, which we show accumulate *in cis* at centromeres. Previous studies by ourselves and others have provided evidence that accumulation of transcripts on chromatin can impair heterochromatin assembly, possibly through increased formation of RNA-DNA hybrids (24,35). Therefore, together, our data suggest that Cpn1 influences the efficiency of heterochromatin assembly by enabling the degradation of heterochromatic transcripts to prevent their detrimental excessive accumulation on chromatin.

Our study revealed the previously uncharacterised protein SPAC12G12.07c as the fission yeast ortholog of human CAPRIN1. Caprin proteins are well conserved in vertebrates, and have also been identified in some other metazoan groups including insects (53); this work confirms that they are also present in fungi. Human CAPRIN1 is implicated in regulating the transport, translation and/or degradation of mRNAs linked to various processes including the innate immune response (54) cell proliferation (47,55) and differentiation (52). However, its best characterised role is in stress granule formation, acting together with partner proteins G3BP1 and USP10. We have confirmed that fission yeast Cpn1 also localises to stress granules, and associates with Nxt3 and Ubp3, which are the orthologs of G3BP1 and USP10, respectively. We also show that Cpn1 is required for granule formation in response to heat shock. This is consistent with studies in human cells that have shown RNA-binding protein G3BP1 to be a key nucleator of stress granules, with CAPRIN1 association acting cooperatively to enhance multivalent interactions and hence promote granule formation, and binding of USP10 (which lacks an RNA-binding domain) conversely ‘capping’ interactions and suppressing granule formation (44,56–58). In line with this, we show here that absence of G3BP1 ortholog Nxt3, but not Usp10 ortholog Ubp3, impairs for granule formation. Superficially, this contrasts with the conclusion from a previous study that Nxt3 is dispensable for granule formation in fission yeast (45). However, the analysis in that study was not quantitative, and we show here that granule formation is reduced but not abolished in *nxt3Δ* cells, indicating that Nxt3, like Cpn1, enhances, but is not absolutely required for, granule formation in fission yeast.

What could be the relationship between stress granule-associated factors and heterochromatin? We have found evidence of interplay between heterochromatin integrity and formation of cytoplasmic RNP granules, since in heterochromatin mutants we observed both increased Pabp-containing granule formation in the absence of stress, and decreased granule formation in response to stress. This suggests that the granules arising in response to heterochromatin deficiency may be formed at the expense of canonical stress granules, suggesting competition for limiting factors. Stress granule formation is typically triggered by translational arrest, which leads to a sudden release of relatively protein-free and unfolded mRNAs from polysomes (59). Interactions between these RNAs and aggregation-prone RNA-binding proteins leads to their phase separation into visible granules. We suggest that in the absence of heterochromatin, accumulating non-coding heterochromatic transcripts may also be bound by some of the same proteins. This would be consistent with our observation that in conditions where these proteins aggregate into stress granules, heterochromatic RNAs can be seen to co-localise. The increased occurrence of Pabp- and Cpn1-containing cytoplasmic granules in heterochromatin-deficient cells in the absence of stress may also reflect the presence of heterochromatic RNPs that occasionally coalesce into visible granules. Although we were not able to detect stable accumulation of heterochromatic RNAs in these granules by RNA-FISH, we suspect that this may be because these granules are associated with RNA turnover, although it is also possible that they arise as a result of altered RNP homeostasis more broadly.

Our observation of crosstalk between heterochromatic silencing and stress granules is reminiscent of a similar phenomenon reported in human cells, where active siRNA-mediated silencing was found to antagonise stress granule formation. In that case, the effect was suggested to be explained by limiting factor Ago2 being required for both processes (60). Here, it is absence of RNAi-directed heterochromatin formation that antagonises stress granule formation, which we suggest could reflect a role for factors such as Cpn1 in mopping up both transcripts released in response to stress, and those accumulating as a result of de-repression of heterochromatin. In simple terms, if RNA-binding proteins are drawn into interactions with accumulated heterochromatic transcripts independent of stress, this could deplete the pool of proteins available to bind translationally arrested RNAs in response to stress, explaining the reduction in stress granule formation we observe in heterochromatin-deficient cells. Given that loss of heterochromatin is widely associated with ageing (61), such alterations in RNP homeostasis could also contribute to broader cellular dysregulation linked to age-related de-repression of heterochromatic domains.

The link between Cpn1 and regulation of heterochromatic transcripts is supported by our observation that absence of Cpn1 functionally impacts the accumulation of pericentromeric RNAs in heterochromatin-deficient cells, causing hyperaccumulation of these RNAs at centromeres that likely explains the observed defect in pericentromeric heterochromatin establishment. This suggests that degradation of heterochromatic transcripts is impaired in *cpn1Δ* cells. Interestingly, human CAPRIN1 was recently shown to promote selective transcript degradation, mediated by Xrn2, a normally nuclear ribonuclease that relocalises to the cytoplasm in a Caprin-dependent manner (52,62). Here we present evidence consistent with possible conservation of this functional connection in fission yeast, since we show that: (1) inactivation of the Xrn2 ortholog, Dhp1, in *clr4Δ* cells results in accumulation of pericentromeric transcripts in the cytoplasm; and (2) this cytoplasmic RNA accumulation is abolished in the absence of Cpn1. This suggests that Cpn1 is required upstream of Dhp1, possibly to facilitate delivery of RNAs for degradation. This would also be consistent with previous studies showing that Dhp1, like Cpn1, is required for efficient heterochromatin establishment, in a manner dependent on its nuclease activity (22,23). In comparison to Cpn1, loss of Dhp1 appears to have a greater impact on heterochromatin maintenance, including at facultative heterochromatin islands and HOODs, consistent with Dhp1 having additional, Cpn1-independent functions in promoting heterochromatin formation. Conversely, nucleases other than Dhp1 may also contribute Cpn1-dependent degradation; these could include, for example, Cpn1 binding partner Nxt3, the human ortholog of which, G3BP1, has been shown to function as an endoribonuclease for degradation of selected RNAs (63).

An outstanding question from our work is whether Cpn1 itself interacts directly with heterochromatic transcripts. Cpn1 is predicted to be an RNA-binding protein, and we found that the potentially RNA-binding RG/RGG-rich region is important for Cpn1 function in promoting efficient heterochromatin establishment. However, we did not detect stable association of Cpn1 with *cen*(*dg*) transcripts by RNA-IP, suggesting that any interaction may be weak/transient. Several lines of evidence support that physical association could occur, despite the superficially distinct localisation of Cpn1 and heterochromatic RNAs to cytosol and nucleus, respectively. Most notably, as referred to above, we have uncovered here the capacity for heterochromatic transcripts to be translocated from the nucleus to the cytoplasm, since in heterochromatin-deficient cells they can be observed to accumulate in cytoplasmic foci in response to stress, and upon suppression of Dhp1-mediated degradation. It may be that detection of these RNAs in the cytoplasm is only possible in conditions where they are aggregated (via their associated RBPs) in stress granules, or their normal rapid degradation otherwise suppressed. Cytosolic localisation of heterochromatic transcripts is not without precedent, since in mouse pancreatic cancer models, aberrantly expressed centromeric satellite repeat RNAs have been seen to accumulate in the cytoplasm (64). In addition, we note that both Cpn1 and Nxt3, although primarily cytoplasmic, have been shown to accumulate in the nucleus following treatment of cells with leptomycin B, which inhibits CRM1-dependent nuclear export (65). This suggests that, like many RNA-binding proteins, they have the capacity to shuttle between the cytoplasm and the nucleus, and so could play a role in nucleocytoplasmic transport. Consistent with this, the human ortholog of Nxt3, G3BP1, has been shown to undergo phosphorylation-dependent nuclear import (63). Taking everything together, the simplest model to explain our findings is that Cpn1 associates transiently with heterochromatic transcripts, in the nucleus and/or cytoplasm, to help target them for degradation (Figure 7A). However, from our data we cannot exclude the possibility that the effect may be mediated by other factors: in the absence of Cpn1 there may be a compensatory shift in the balance of binding of other RBPs, resulting in reduced availability of one or more other factors that would normally associate with heterochromatic transcripts to promote their degradation (Figure 7B). Either way, our results suggest an important connection between heterochromatin assembly and RNP homeostasis.

**Figure 7.**
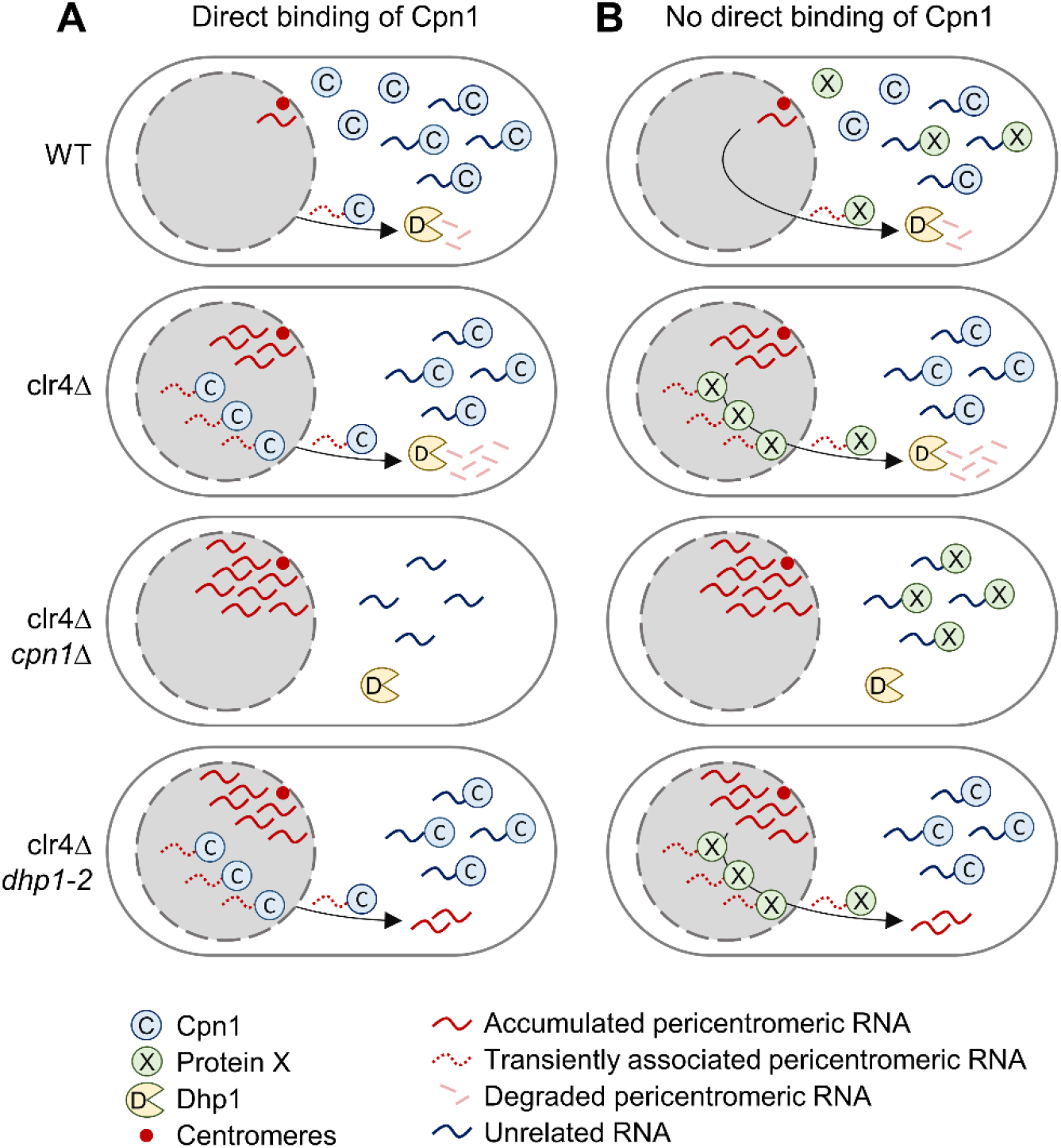
Models for impact of Cpn1 on pericentromeric transcript regulation. (A) Cpn1 transiently binds pericentromeric RNAs to target them for Dhp1-dependent degradation. In comparison to wild-type cells (top), in *clr4Δ* cells there is increased pericentromeric transcript accumulation leading to greater Cpn1 association. Absence of Cpn1 directly leads to reduced targeting for degradation and hence further nuclear RNA accumulation; suppression of Dhp1-mediated degradation leads to RNA accumulation in the nucleus and cytoplasm. (B) Cpn1 does not directly bind pericentromeric transcripts and effects are mediated via altered RNP homeostasis. Protein X directly binds pericentromeric RNAs to target them for Dhp1-dependent degradation. However, in the absence of Cpn1, Protein X is engaged in binding other RNAs at the expense of pericentromeric transcripts, leading to reduced degradation. In both models, increased binding to pericentromeric transcripts in *clr4Δ* cells compared to wild-type reduces the free RBP pool available for stress granule formation in response to stress. RBP binding and Dhp1-mediated degradation may occur in the nucleus and/or cytoplasm.

Human CAPRIN1 is increasingly being recognised as a protein of interest in several clinical contexts. For example, it contributes to antiviral responses, and several viruses, including Zika virus, have been found to hijack CAPRIN1 to inhibit stress granule formation and thus promote viral replication (66). Moreover, CAPRIN1 also plays important roles in brain development, regulating the transport and translation of mRNAs required for synaptic plasticity. Deficiency of CAPRIN1 has been linked to autism spectrum disorders (67) and long-term memory impairment (68), while a specific missense mutation (P512L) has recently been implicated in early-onset ataxia and neurodegeneration (69). In addition, changes in CAPRIN1 expression and localisation have also been linked to cancer; in particular, the recent detection of CAPRIN1 on the cell membrane surface in many solid cancers, but not normal tissues, makes it an attractive novel target for cancer therapeutics (70). Despite their potential importance, the roles of CAPRIN1 outside of stress granules remain poorly understood; the identification of a Caprin family protein in fission yeast opens new opportunities to dissect the molecular mechanisms underpinning the cellular functions of these proteins, as well as unveiling a novel connection between RNP homeostasis and heterochromatin assembly.

## Materials and Methods

### Yeast strains and methods

Strains used in this study are listed in Table S1. Standard procedures were used for growth and genetic manipulations. Cultures were grown at 32°C unless otherwise stated, in either rich medium (YES) or minimal media (PMG). Genomic integrations for gene deletion and epitope-tagging were achieved by homologous recombination using PCR-based modules consisting of a resistance cassette flanked by 80 bp sequences homologous to the target locus. To generate the *cpn1^R6A^* mutant, the six arginines in the RGG motif were mutated to alanine by CRISPR/Cas9 editing as previously described (71). Genomic modifications were verified by sequencing. Overexpression plasmids pSP102-Sir2, pSP102-Clr3 and pSP102-Sir2-Clr3 were described previously (72). The minichromosome establishment assay was carried out as described previously (32). For cross-based establishment assays, random spore analysis was employed. Strains bearing different gene deletions marked with *ura4^+^* were crossed, and the products plated onto selective media containing 1g/L 5-FOA and low (1/10^th^) adenine to directly select for desired recombinant genotypes (and against parental genotypes) prior to counting of red/white colonies.

### Live cell Imaging

Visualisation of fluorescently-tagged proteins in living cells was performed on log phase cultures grown at 32°C in PMG and, where indicated, pre-treated as follows: heat shock, 20 min incubation at 42°C; glucose starvation, 20 min incubation in medium lacking glucose. Images were acquired using a Nikon Ti2 inverted microscope equipped with a 100x 1.49 NA Apo TIRF objective and a Teledyne Photometrics Prime 95B camera. Images were acquired with NIS-elements (version 5.1), with 25 z-stacks taken at 0.2µm intervals, deconvolved with AutoQuant X3 software and subsequently exported to ImageJ for analysis.

### smRNA FISH

smRNA FISH was performed essentially as described previously (73). Briefly, 2×10^8^ cells were fixed with 4% paraformaldehyde for 30 min. Cells were washed with Buffer B (1.2 M sorbitol, 0.1M potassium phosphate buffer pH 7.5), resuspended in spheroplast buffer (1.2 M sorbitol, 0.1 M potassium phosphate, 20 mM vanadyl ribonuclease complex [NEB, S1402S], 20 μM beta-mercaptoethanol) and digested with 0.002% 100 T zymolyase (US Biological, Z1005) for approximately 45–75 min. Cells were washed with Buffer B, resuspended in 1 ml of 0.01% Triton X-100 in 1x PBS for 20 min, washed again with Buffer B, and resuspended in 10% formamide/2x SSC buffer. For hybridization of probes, approximately 20–25ng of CAL Fluor red 610 probes targeting *cen(dg)* repeat sequences were mixed with 2 μl each of yeast tRNA (Life Technologies) and Salmon sperm DNA (Life Technologies) per reaction. Sequences of the *cen(dg)* probes are given in Table S3. The probe solution was mixed with Buffer F (20% formamide, 10 mM sodium-phosphate buffer pH 7.2; 45 μl per reaction), heated at 95°C for 3 min, and allowed to cool to room temperature before mixing with Buffer H (4x SSC, 4 mg/ml acetylated BSA, 20 mM vanadyl ribonuclease complex; 50 μl per reaction). Digested cell samples were divided into two reactions, each of which was resuspended in 100 μl of this hybridization solution and incubated at 37°C overnight. Cells were subsequently washed sequentially with: 10% formamide/2x SSC; 0.1% Triton X-100/2x SSC; 2x SSC; 1x SSC; and 1x PBS. Cell pellets were mixed with VECTASHIELD Vibrance® Antifade Mounting Medium with DAPI (Vector Laboratories) and mounted on poly-L-Lysine slides with 1.5mm glass coverslips. Imaging was performed using a Nikon Ti2 inverted microscope equipped with a 100x 1.49 NA Apo TIRF objective and a Teledyne Photometrics Prime 95B camera. Images were acquired with NIS-elements (version 5.1), with 25 z-stacks taken at 0.2µm intervals, further deconvolved with AutoQuant X3 software and subsequently exported to ImageJ for analysis.

### RT-qPCR

Total RNA was extracted from 1 × 10^7^ cells in exponential growth phase using the Masterpure Yeast RNA Purification Kit (Epicentre) according to the manufacturer’s instructions. 1 μg of total RNA was treated with TURBO DNase (Ambion) for 1 h at 37°C, then reverse transcribed using random hexamers (Roche) and Superscript III reverse transcriptase (Invitrogen) according to the manufacturer’s instructions. cDNA was quantified by qPCR using LightCycler 480 SYBR Green (Roche) and primers listed in Supplementary Table S2. In all cases, histograms represent three biological replicates and error bars represent one S.D.

### ChIP

ChIP was performed essentially as described previously (74). Briefly, 2.5 × 10^8^ cells per IP were fixed in 1% formaldehyde for 15 min at room temperature. Cells were lysed using a bead beater (Biospec products, 2 x 2 min) and sonicated using a Bioruptor (Diagenode) for a total of 20 min (30 s on/30 s off on ‘high’ power). Immunoprecipitation was then performed overnight at 4°C, using 1 μl per IP of monoclonal anti-H3K9me2 (5.1.1 (75)). Immunoprecipitated DNA was recovered using Chelex-100 resin (BioRad), and quantified by qPCR using LightCycler 480 SYBR Green (Roche) and primers listed in Supplementary Table S2. Relative enrichments were calculated as the ratio of product of interest to control product (*act1^+^*) in IP over input. In all cases, histograms represent three biological replicates and error bars represent one S.D.

### Immunoprecipitation

For IP-western, cultures were grown to mid-log phase in YES and 3 x 10^8^ cells were harvested, washed and resuspended in lysis buffer (50 mM Hepes pH 7.5, 150 mM NaCl, 5 mM EDTA, 0.1% NP-40, 1 mM PMSF, 1× yeast protease inhibitor cocktail [Sigma]). Cells were lysed by using a bead beater (Biospec products, 2 x 2 min), DTT was added to a final concentration of 500 μM, and the supernatant was recovered by 2 x 15 min centrifugation at 17,000 rcf at 4°C. Extracts were pre-cleared by incubation with pre-equilibrated protein G agarose for 1 hour at 4°C, and then incubated with pre-equilibrated protein G agarose plus anti-FLAG M2 (F3165, Merck, 1 μl/IP), or 5 μl/IP anti-GFP (A-11122, ThermoFisher, 5 ul/IP), for 3 hours at 4°C. The beads were washed three times with lysis buffer, and proteins eluted in 2x SDS sample buffer (50mM Tris-HCl pH 6.8, 2mM EDTA, 10% glycerol, 2% SDS, 2% β-Mercaptoethanol, 0.03% Bromophenol Blue) by boiling for 5 min. For analysis, proteins were separated by SDS-PAGE and transferred onto 0.45μm pore Protran nitrocellulose membrane (GE Healthcare) using semi-dry transfer apparatus (Hoeffer). Membranes were probed with mouse anti-FLAG M2 (F3165, Merck, 1:1000 dilution) or rabbit anti-GFP (A-11122, ThermoFisher, 1:1000 dilution). Secondary antibodies were IRDye® conjugated anti-mouse or anti-rabbit (926-32210 or 926-68171, Li-Cor, 1:20,000 dilution) and imaging was performed using the Odyssey® CLx Imaging System (LI-COR Biosciences).

Immunoaffinity purifications for mass spectrometry analysis were performed essentially as described previously (76). Briefly, 500 ml cultures grown to a cell density of 10^8^ cells/ml in 4× concentrated YES media were milled in solid phase, and the cell powder solubilised in 10ml lysis buffer (50 mM HEPES–NaOH pH 7.5, 150 mM KCl, 0.1% NP-40, 0.2 mM PMSF, 0.2mM benzamidine, 1× EDTA-free protease inhibitor cocktail [Roche]). Following clarification by two rounds of centrifugation, immunoprecipitations performed by incubating with 8 μl Protein G Dynabeads (Novex) and 12 μl anti-FLAG antibody (F3165, Merck) per sample for 90 min at 4°C. The immunoprecipitated material was washed with lysis buffer, incubated in lysis buffer containing 2 mM MgCl_2_ and 500 U of Benzonase nuclease (Novagen) for 15 min at 4°C, washed again with lysis buffer, and then eluted by incubation with 50 μl of 0.1% Rapigest (Waters) in 50 mM Tris-HCl pH 8.0 for 10 min at 50°C prior to LC-MS/MS analysis.

RNA-IPs were performed as above but with modified lysis buffer (50 mM HEPES–NaOH pH 7.5, 150 mM NaCl, 1mM MgCl_2_, 0.1% NP-40, 5 mM DTT, 0.5 mM PMSF, 1× EDTA-free protease inhibitor cocktail [Roche], 0.2U/μl RNAsin Ribonuclease Inhibitor [Promega]), and excluding Benzonase treatment. After initial washes, immunoprecipitated material was resuspended in RNA extraction buffer (25 mM Tris–HCl pH 7.5, 5 mM EDTA pH 8, 50 mM NaCl, 0.5% SDS) with 200 ng/ml proteinase K and incubated at 37°C for 2 h. RNA was extracted with phenol:chloroform, precipitated with ethanol and 1 μl glycogen, and analysed by RT-qPCR.

### Mass spectrometry

IP eluate was reduced with 25 mM DTT at 80°C for 1 min, then denatured by addition of urea to 8M. Sample was applied to a Vivacon 30k MWCO spin filter (Sartorius, VN01H21) and centrifuged at 12,500 g for 15 min. Protein retained on the column was then alkylated with 100 μl of 50 μM iodoacetamide (IAA) in buffer A (8 M urea, 100 mM Tris pH 8.0) in the dark at RT for 20 min. The column was then centrifuged as before, and washed with 100 μl buffer A, then with 2 x 100 μl volumes of 50 mM ammonium bicarbonate (ABC). 3 μg/μl trypsin (Pierce, 90057) in 0.5 mM ABC was applied to the column and incubated at 37° for 16 hr. Digested peptides were then spun through the filter, acidified with trifluoroacetic acid (TFA) to pH <= 3, loaded onto manually-prepared and equilibrated C_18_ reverse-phase resin stage tips (Sigma, 66883-U) (77), washed with 100 μl 0.1% TFA, and stored at -20°C.

For MS analysis, peptides were eluted in 40 μl of 80% acetonitrile in 0.1% TFA and concentrated down to 2 μl by vacuum centrifugation (Concentrator 5301, Eppendorf). The peptide sample was then prepared for LC-MS/MS analysis by diluting it to 5 μl by 0.1% TFA. LC-MS-analyses were performed on an Orbitrap Exploris 480™ Mass Spectrometer (Thermo Scientific) coupled on-line, to an Ultimate 3000 RSLCnano Systems (Dionex, Thermo Fisher Scientific). Peptides were separated on a 50 cm EASY-Spray column (Thermo Scientific), which was assembled on an EASY-Spray source (Thermo Scientific) and operated at 50°C. Mobile phase A consisted of 0.1% formic acid in LC-MS grade water and mobile phase B consisted of 80% acetonitrile and 0.1% formic acid. Peptides were loaded onto the column at a flow rate of 0.3 μl per min and eluted at a flow rate of 0.25 μl per min according to the following gradient: 2 to 40% mobile phase B in 150 min, and then to 95% in 11 min. Mobile phase B was retained at 95% for 5 min and returned back to 2% a minute after until the end of the run (190 min). FTMS spectra were recorded at 120,000 resolution (scan range 350-1500 m/z) with an ion target of 3.0×10^6^ and maximum injection time of 50 ms. For the MS2 the resolution was set at 15,000 with ion target of 8.0×10^4^ and HCD fragmentation (78) with normalized collision energy of 30. The isolation window in the quadrupole was 1.4 Thomson. Only ions with charge between 2 and 6 were selected for MS2.

The MaxQuant software platform (79) version 1.6.1.0 was used to process the raw files and search was conducted against the *Schizosaccharomyces pombe* complete/reference proteome (Pombase, released in July 2017), using the Andromeda search engine (80). For the first search, peptide tolerance was set to 20 ppm while for the main search peptide tolerance was set to 4.5 pm. Isotope mass tolerance was 2 ppm and maximum charge to 7. Digestion mode was set to specific with trypsin allowing a maximum of two missed cleavages. Carbamidomethylation of cysteine was set as fixed modification. Oxidation of methionine was set as variable modification. Absolute protein quantification was performed as described previously (81). Peptide and protein identifications were filtered to 1% FDR.

## Data Availability

Mass spectrometry data have been deposited to the ProteomeXchange Consortium via the PRIDE repository with the dataset identifier PXD050937.

## Acknowledgements

We thank Robin Allshire for sharing Sir2 and Clr3 over-expression plasmids and GFP-Cnp1 strain, Ke Zhang for the *dhp1-2* strain, and Takeshi Urano for provision of anti-H3K9me2 antibody. We are also grateful to Dave Kelly and Toni McHugh for assistance with imaging analysis, Silke Hauf for advice on smRNA-FISH, Ana Arsenijevic for guidance on IP-MS sample prep and data analysis, and Elliott Chapman for helpful discussions and critical reading of the manuscript. Imaging was performed in Centre Optical Instrumentation Laboratory (COIL), supported by a Core Grant (203149) to the Wellcome Centre for Cell Biology at the University of Edinburgh.

## Supplementary Information

**Figure S1.**
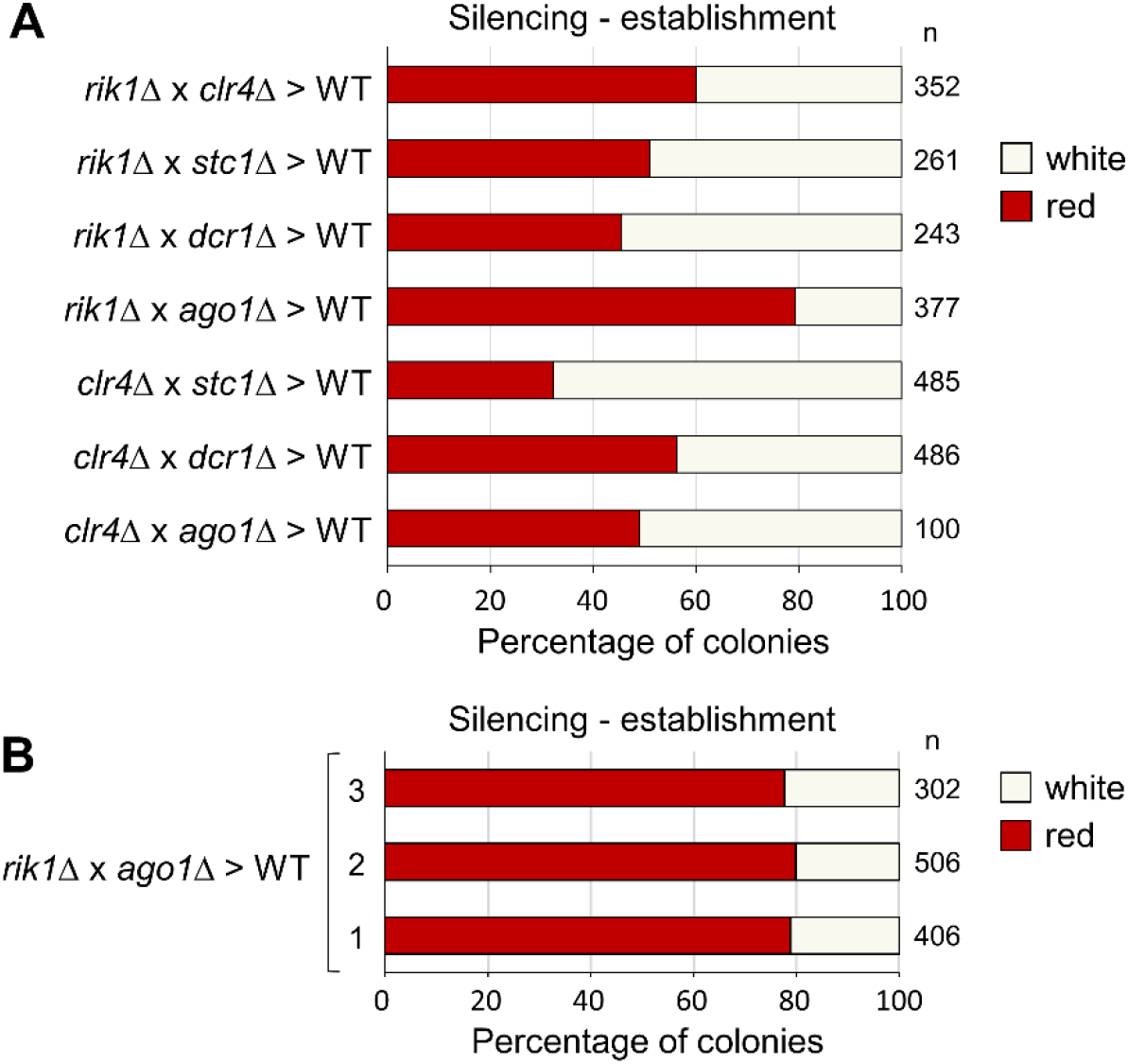
Efficiency of heterochromatin re-establishment varies with genetic background, but is consistent for the same background. (A) Proportions of red (*cen1:ade6^+^* silenced) versus white (*cen1:ade6^+^* expressed) colonies in the wild-type progeny of crosses between the indicated parental strains, based on analysis of *n* colonies. (B) Proportions of red (*cen1:ade6^+^* silenced) versus white (*cen1:ade6^+^* expressed) colonies in the wild-type progeny of crosses between three independent *rik1Δ* and *ago1Δ* strains, based on analysis of *n* colonies.

**Figure S2.**
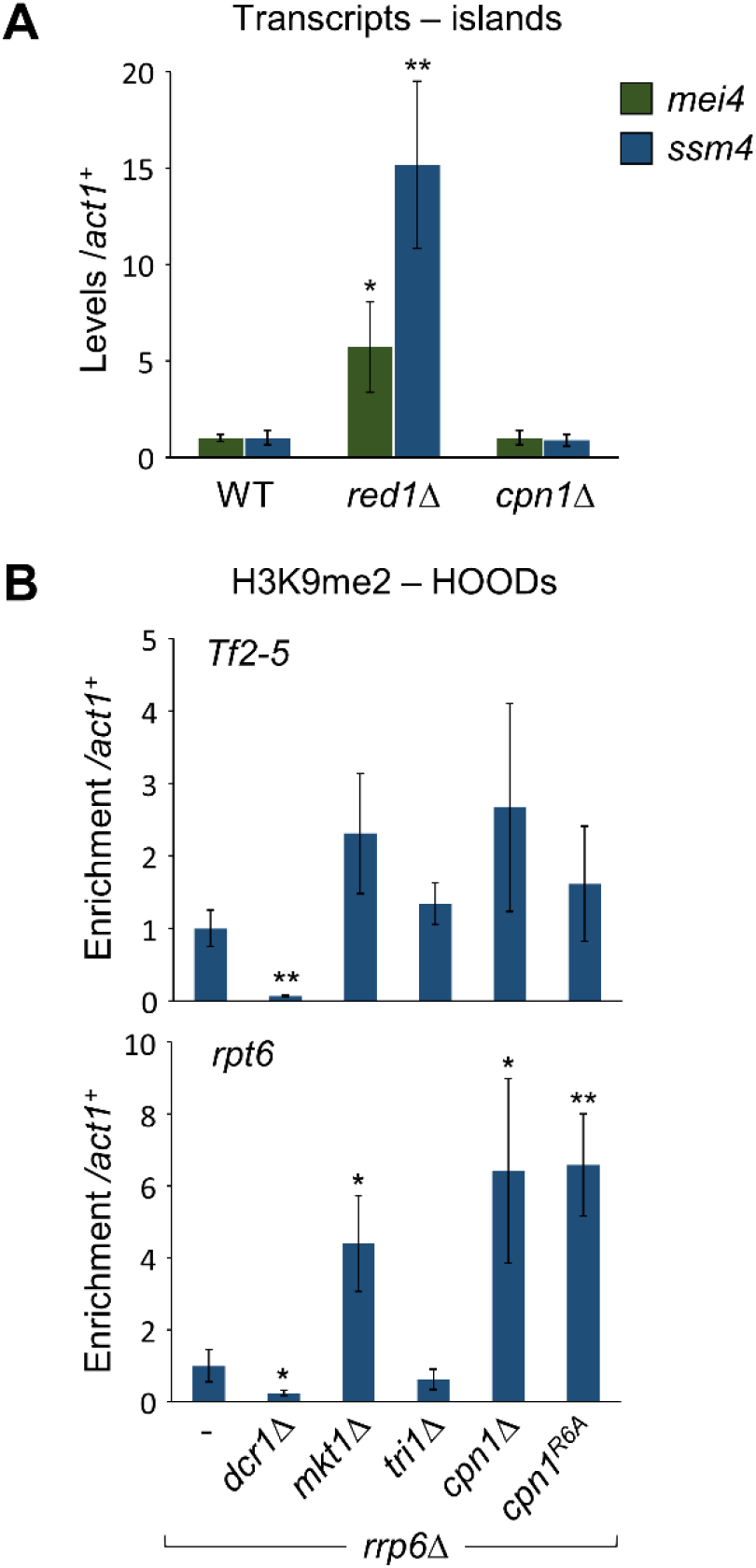
Cpn1 is not required for silencing at heterochromatin islands or most HOODs. (A) RT-qPCR analysis of indicated transcript levels relative to *act1^+^*, normalized to wild-type. (B) ChIP-qPCR analysis of H3K9me2 levels at indicated HOODs, relative to *act1^+^*, normalised to wild-type. Data are averages of three biological replicates, error bars represent one SD. Relative to wild-type, asterisks denote p ≤ 0.05 (*), or p ≤ 0.01 (**), from Student’s t-test analysis.

**Figure S3.**
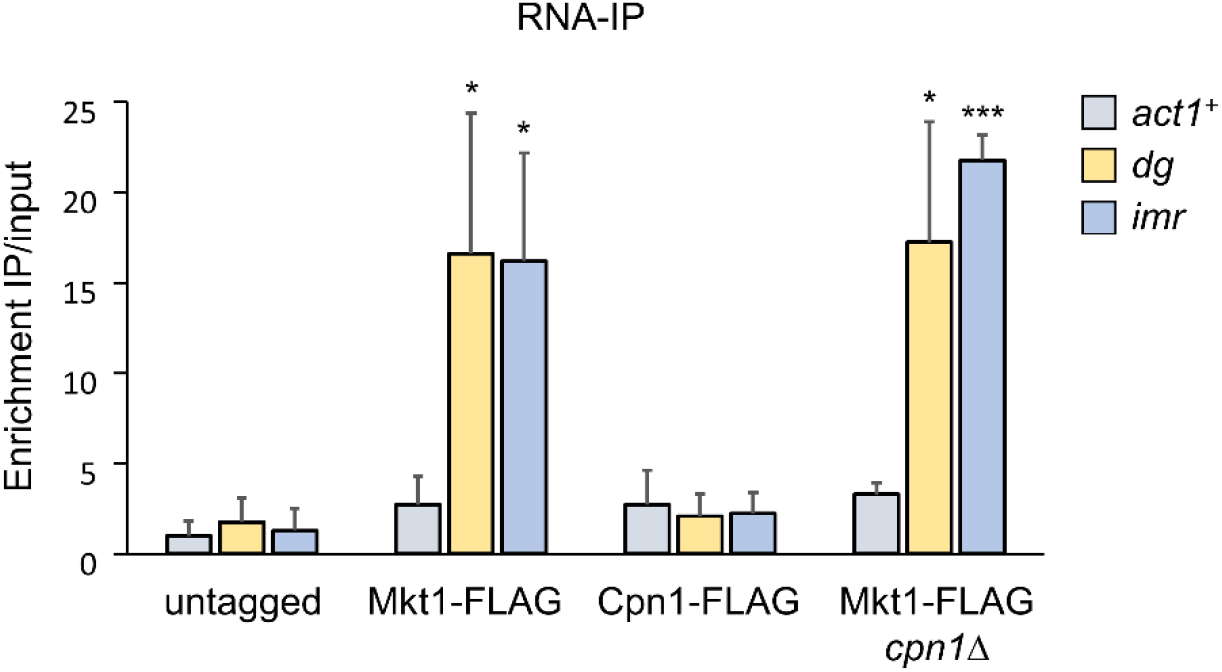
Mkt1 but not Cpn1 interacts with centromeric transcripts. RNA-immunoprecipitation (RNA-IP) analysis of transcripts associated with FLAG-tagged Cpn1 or Mkt1 (wild-type or *cpn1Δ* backgrounds), under native conditions. IP enrichments are shown relative to input. Data are averages of three biological replicates, error bars represent one SD. Relative to untagged control, asterisks denote p ≤ 0.05 (*), or p ≤ 0.001 (***), from Student’s t-test analysis.

**Figure S4.**
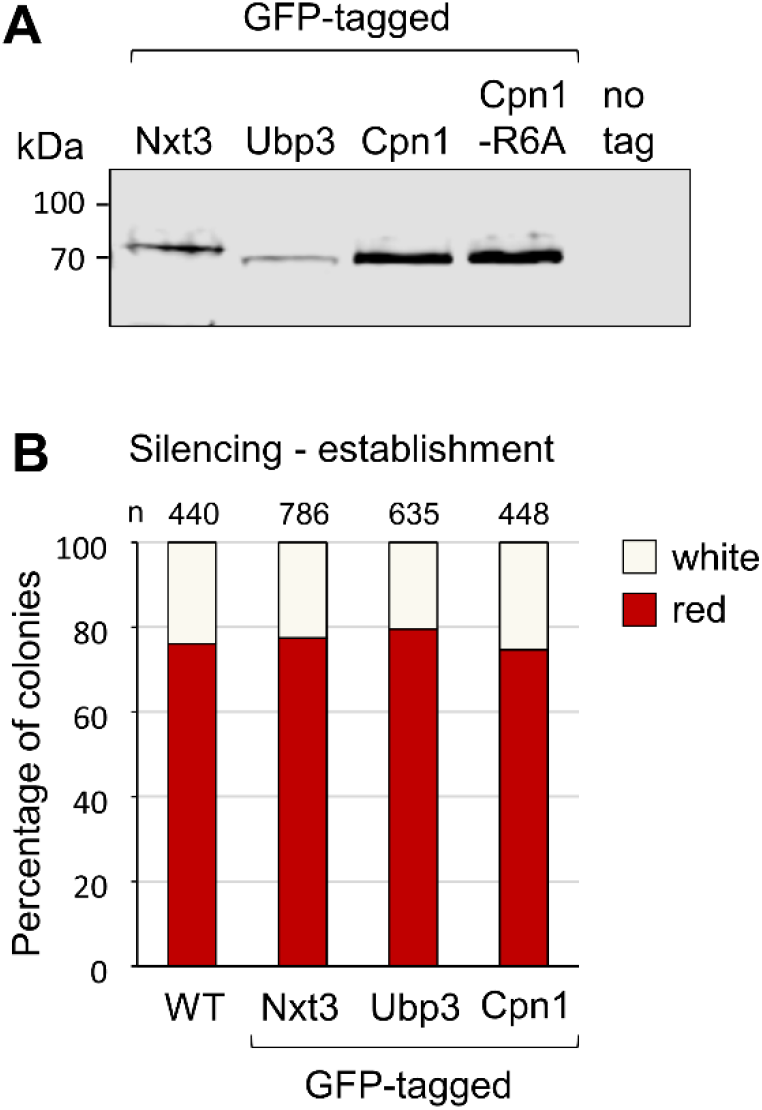
GFP-tagged Cpn1, Nxt3 and Ubp3 are stable and functional. (A) Western blot analysis of affinity-purified GFP-tagged Nxt3, Ubp3, Cpn1, and Cpn1^R6A^. (B) Proportions of red (*cen1:ade6^+^* silenced) versus white (*cen1:ade6^+^* expressed) colonies in the otherwise wild-type progeny of *rik1Δ* x *ago1Δ* crosses performed in the indicated genetic backgrounds, based on analysis of *n* colonies.

**Figure S5.**
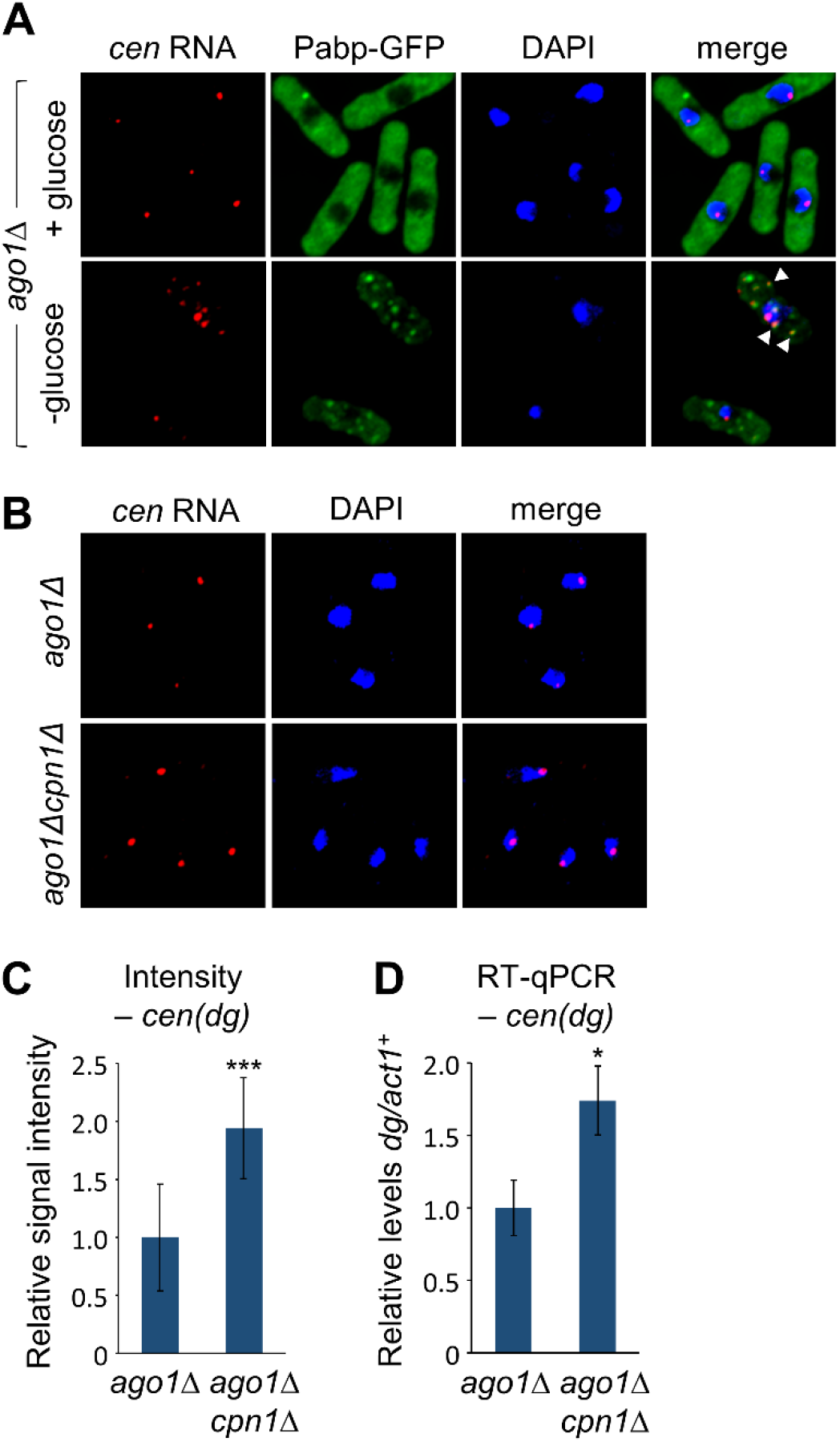
Absence of Cpn1 results in increased accumulation of pericentromeric transcripts at centromeres in *ago1Δ* cells. (A) Representative images from simultaneous analysis of *cen(dg)* RNA by smRNA-FISH, and Pabp-GFP, in *ago1Δ* cells either unstressed (+ glucose) or stressed by 20 min of glucose starvation (-glucose). Arrow heads highlight examples of co-localisation. (B) Representative images from smRNA-FISH analysis of *cen(dg)* RNA in *ago1Δ* and *ago1Δ cpn1Δ* cells. (B) Quantification of the mean signal intensity for nuclear *cen(dg)* RNA foci in *ago1Δ cpn1Δ* relative to *ago1Δ* cells, from the smRNA-FISH analysis shown in B. (D) RT-qPCR analysis of total cellular levels of *cen*(*dg*) RNA relative to *act1^+^*, in *ago1Δ cpn1Δ* relative to *ago1Δ* cells. RT-qPCR data are averages of three biological replicates. In all cases, error bars represent one SD, and asterisks denote p ≤ 0.05 (*), or p ≤ 0.001 (***), from Student’s t-test analysis.

**Table S1.** Yeast strains.

| Strain | Genotype | Figure |
| --- | --- | --- |
| 7 | *h+ otr1R::ade6+ ade6-210 leu1-32 ura4-D18* | 1,2,3,S2,S3 |
| 131 | *h- dcr1::NatMX6 ade6-210 leu1-32 ura4-D18 arg3-D4 his3D1* | 3 |
| 140 | *h- sir2::NatMX6 ade6-210 otrR::ade6+ ura4-D18* | S4 |
| 1621 | *h+ rrp6::KanMX6 ade6-210 ura4-D18 leu1-32* | 2,S2 |
| 1952 | *h- ago1::ura4+ otr1R::ade6+ ade6-210 ura4-D18 leu1-32* | 2 |
| 2233 | *rrp6::KanMX6 mkt1::NatMX6 otr1R::ade6+ ade6-210 leu1-32 ura4-D18* | 2,S2 |
| 2505 | *rrp6::KanMX6 dcr1::NatMX6 otr1R::ade6+ ade6-210 leu1-32 ura4-DSE* | 2,S2 |
| 4471 | *h90 clr4::leu2+ cc2::his3+ ade6-704-NatMX6 ura4-DSE/D18 leu1-32 his3-D1* | 2 |
| 4472 | *h+ cc2::his3+ ade6-704-HygMX6 ura4-DSE/D18 leu1-32 his3-D1 arg3-D4* | 2 |
| 4512 | *h- sir2::ura4+ Mkt1-Flag-NatMX6 otr1R::ade6+ ade6-210 leu1-32 ura4-D18* | S4 |
| 4578 | *h- rik1::HygMX6:ura4+ otr1R::ade6+ ade6-210 leu1-32 ura4-D18 his3-D1* | 1,2,3,S1,S3 |
| 4658 | *h+ dcr1::NatMX6:ura4+ otr1R::ade6+ ade6-210 leu1-32 ura4-D18* | S1 |
| 5135 | *h+ ago1::NatMX6:ura4+ otr1R::ade6+ ade6-210 leu1-32 ura4-D18* | 1,2,3,S1,S3 |
| 5288 | *h+ stc1::NatMX6:ura4+ otr1R::ade6+ ade6-210 leu1-32 ura4-D18* | S1 |
| 6407 | *red1::KanMX6 otr1R::ade6+ ade6-210 leu1-32 ura4-D18 arg3-D4* | S2 |
| 6601 | *h90 ago1::NatMX6:ura4+ lys1::leu1+ otr1R::ade6+ ade6-210 leu1-32 ura4-D18* | 1,S1 |
| 6605 | *h- rik1::HygMX6:ura4+ lys1::leu1+ otr1R::ade6+ ade6-210 leu1-32 ura4-D18* | 1,S1 |
| 6633 | *h+ rik1::HygMX6:ura4+ ago1::NatMX6:ura4+ lys1::leu1+ otr1R::ade6+ ade6-210 leu1-32 ura4-D18* | 1 |
| 6639 | *h- ago1::NatMX6:ura4+ otr1R::ade6+ ade6-DN/N leu1-32 ura4-D18 his3-D1* | S1 |
| 6751 | *h- ago1::NatMX6:ura4+ mkt1::KanMX6 otr1R::ade6+ ade6-210 leu1-32 ura4-D18* | 1 |
| 6759 | *h+ ago1::NatMX6:ura4+ tri1::KanMX6 otr1R::ade6+ ade6-210 leu1-32 ura4-D18* | 1 |
| 6765 | *h+ rik1::HygMX6:ura4+ mkt1::KanMX6 otr1R::ade6+ ade6-210 leu1-32 ura4-D18* | 1 |
| 6798 | *h- rik1::HygMX6:ura4+ tri1::KanMX6 otr1R::ade6+ ade6-210 leu1-32 ura4-D18* | 1 |
| 6928 | *h+ otr1R::ade6+ lys1::leu1+ ura4+ ade6-210 leu1-32* | 1 |
| 6974 | *h- clr4::HygMX6:ura4+ otr1R::ade6+ ade6-210 leu1-32 ura4-D18 his3-D1* | 1,2,3,6,S1 |
| 7089 | *rik1::HygMX6:ura4+ otr1R::ade6+ ade6-DN/N leu1-32 ura4-D18 his3-D1* | S1 |
| 7120 | *h+ PSP102-Sir2 leu2+ rik1::HygMX6:ura4+ otr1R::ade6+ ade6-210 leu1-32 ura4-D18 his3-D1* | 1 |
| 7123 | *h- PSP102-Sir2 leu2+ ago1::NatMX6:ura4+ otr1R::ade6+ ade6-210 leu1-32 ura4-D18 his3-D1* | 1 |
| 7124 | *h+ PSP102-Clr3 leu2+ rik1::HygMX6:ura4+ otr1R:ade6+ ade6-210 leu1-32 ura4-D18 his3-D1* | 1 |
| 7129 | *h- PSP102-Clr3 leu2+ ago1::NatMX6:ura4+ otr1R:ade6+ ade6-210 leu1-32 ura4-D18 his3-D1* | 1 |
| 7130 | *h+ PSP102-Sir2&Clr3 leu2+ rik1::HygMX6:ura4+ otr1R:ade6+ ade6-210 leu1-32 ura4-D18 his3-D1* | 1 |
| 7133 | *h- PSP102-Sir2&Clr3 leu2+ ago1::NatMX6:ura4+ otr1R:ade6+ ade6-210 leu1-32 ura4-D18 his3-D1* | 1 |
| 7221 | *h- gcn5::KanMX6 rik1::HygMX6:ura4+ otr1R:ade6+ ade6-210 leu1-32 ura4-D18* | 1 |
| 7225 | *h+ gcn5::KanMX6 ago1::NatMX6:ura4+ otr1R:ade6+ ade6-210 leu1-32 ura4-D18* | 1 |
| 7577 | *h+ cpn1::HygMX6 otr1R:ade6+ ade6-210 leu1-32 ura4-D18* | 2,3,S2 |
| 7669 | *h+ cpn1::KanMX6 rik1::HygMX6:ura4+ otr1R:ade6+ ade6-210 leu1-32 ura4-D18 his3-D1* | 2,3 |
| 7706 | *h- cpn1::KanMX6 ago1::NatMX6:ura4+ otr1R:ade6+ ade6-210 leu1-32 ura4-D18* | 2,3 |
| 7707 | *h+ cpn1::KanMX6 ago1::NatMX6:ura4+ otr1R:ade6+ ade6-210 leu1-32 ura4-D18* | 2 |
| 7805 | *h- ago1::ura4+ otr1R:ade6+ lys1::NatMX6 ade6-210-HygMX6 mat1_m-cyhS smt0 CycR leu1-32 ura4-D18* | S1 |
| 7905 | *h+ rik1::ura4+ otr1R:ade6+ lys1::NatMX6 ade6-210-HygMX6 ura4-D18 leu1-32* | S1 |
| 8457 | *rrp6::KanMX6 cpn1::Hyg otr1R:ade6+ ade6-210 ura4-D18 leu1-32* | 2,S2 |
| 8543 | *h+ Cpn1-GFP-KanMX6 otr1R:ade6+ ade6-210 leu1-32 ura4-D18* | 6,S3 |
| 8603 | *h+ cpn1<sup>R6A</sup> otr1R:ade6+ ade6-210 leu1-32 ura4-D18* | 2,3 |
| 8650 | *rrp6::KanMX6 tri1::leu1+ otr1R:ade6+ ade6-210 ura4-D18 leu1-32* | 2,S2 |
| 8725 | *h+ cpn1<sup>R6A</sup> ago1::NatMX6:ura4+ otr1R:ade6+ ade6-210 leu1-32 ura4-D18* | 2 |
| 8776 | *rrp6::KanMX6 cpn1<sup>R6A</sup> otr1R:ade6+ ade6-210 ura4-D18 leu1-32* | 2,S2 |
| 8800 | *cpn1::KanMX6 cc2::his3+ ade6-704-HygMX6 ura4-DSE/D18 leu1-32* | 2 |
| 8314 | *h+ Cpn1-Flag-NatMX6 otr1R:ade6+ ade6-210 leu1-32 ura4-D18* | 3 |
| 8681 | *h- cpn1<sup>R6A</sup> rik1::HygMX6:ura4+ otr1R:ade6+ ade6-210 leu1-32 ura4-D18* | 3 |
| 8725 | *h+ cpn1<sup>R6A</sup> ago1::NatMX6:ura4+ otr1R:ade6+ ade6-210 leu1-32 ura4-D18* | 3 |
| 8785 | *Cpn1-GFP-KanMX6 rik1::HygMX6:ura4+ otr1R:ade6+ ade6-210 leu1-32 ura4-D18* | S3 |
| 8787 | *Cpn1-GFP-KanMX6 ago1::NatMX6:ura4+ otr1R:ade6+ ade6-210 leu1-32 ura4-D18* | S3 |
| 8789 | *h+ Nxt3-GFP-KanMX6 otr1R:ade6+ ade6-210 leu1-32 ura4-D18* | S3 |
| 8798 | *cpn1::NatMX6 clr4::HygMX6:ura4+ otr1R:ade6+ ade6-210 leu1-32 ura4-D18* | 6 |
| 8804 | *h+ Ubp3-GFP-KanMX6 otr1R:ade6+ ade6-210 leu1-32 ura4-D18* | S3 |
| 8834 | *h+ Cpn1<sup>R6A</sup>-GFP-KanMX6 otr1R:ade6+ ade6-210 leu1-32 ura4-D18* | S3 |
| 8835 | *h+ ubp3::HA-KanMX6 otr1R:ade6+ ade6-210 leu1-32 ura4-D18* | 3 |
| 8854 | *h+ Cpn1<sup>R6A</sup>-Flag-NatMX6 otr1R:ade6+ ade6-210 leu1-32 ura4-D18* | 3 |
| 8869 | *clr4::HygMX6:ura4+ Cpn1-GFP-KanMX6 otr1R:ade6+ ade6-210 leu1-32 ura4-D18* | 6 |
| 8872 | *Nxt3-GFP-KanMX6 rik1::HygMX6:ura4+ otr1R:ade6+ ade6-210 leu1-32 ura4-D18 his3-D1* | S3 |
| 8874 | *Ubp3-GFP-KanMX6 rik1::HygMX6:ura4+ otr1R:ade6+ ade6-210 leu1-32 ura4-D18 his3-D1* | S3 |
| 8878 | *Nxt3-GFP-KanMX6 ago1::NatMX6:ura4+ otr1R:ade6+ ade6-210 leu1-32 ura4-D18* | S3 |
| 8880 | *Ubp3-GFP-KanMX6 ago1::NatMX6:ura4+ otr1R:ade6+ ade6-210 leu1-32 ura4-D18* | S3 |
| 8928 | *ubp3::HA-KanMX6 rik1::HygMX6:ura4+ otr1R::ade6+ ade6-210 leu1-32 ura4-D18* | 3 |
| 8930 | *ubp3::HA-KanMX6 ago1::NatMX6:ura4+ otr1R::ade6+ ade6-210 leu1-32 ura4-D18* | 3 |
| 8948 | *h- nxt3::HA-KanMX6 otr1R:ade6+ ade6-210 leu1-32 ura4-D18* | 3 |
| 9031 | *nxt3::HA-KanMX6 rik1::HygMX6:ura4+ otr1R:ade6+ ade6-210 leu1-32 ura4-D18* | 3 |
| 9033 | *nxt3::HA-KanMX6 ago1::NatMX6:ura4+ otr1R:ade6+ ade6-210 leu1-32 ura4-D18* | 3 |
| 9144 | *h+ Cpn1-GFP-KanMX6 Pabp-mRFP-HygMX6 leu1-32 ura4-D18* | 4,5 |
| 9145 | *h+ Cpn1<sup>R6A</sup>-GFP-KanMX6 Pabp-mRFP-HygMX6 leu1-32 ura4-D18* | 4 |
| 9147 | *h+ cpn1::NatMX6 Pabp-mRFP-HygMX6 leu1-32 ura4-D18* | 4,5 |
| 9151 | *h+ Nxt3-GFP-KanMX6 Pabp-mRFP-HygMX6 leu1-32 ura4-D18* | 4 |
| 9152 | *h+ Ubp3-GFP-KanMX6 Pabp-mRFP-HygMX6 leu1-32 ura4-D18* | 4 |
| 9225 | *mkt1::NatMX6 Cpn1-GFP-KanMX6 Pabp-mRFP-HygMX6 leu1-32 ura4-D18* | 4,5 |
| 9226 | *ubp3::NatMX6 Cpn1-GFP-KanMX6 Pabp-mRFP-HygMX6 leu1-32 ura4-D18* | 4 |
| 9232 | *cpn1<sup>R6A</sup> Pabp-mRFP-HygMX6 leu1-32 ura4-D18* | 4 |
| 9263 | *clr4::NatMX6 Cpn1-GFP-KanMX6 Pabp-mRFP-HygMX6 leu1-32 ura4-D18* | 5 |
| 9297 | *nxt3::NatMX6 Cpn1-GFP-KanMX6 Pabp-mRFP-HygMX6 leu1-32 ura4-D18* | 4 |
| 9306 | *h90 ago1::ura4+ Cpn1-GFP-KanMX6 Pabp-mRFP-HygMX6 leu1-32 ura4-D18* | 5 |
| 9327 | *Cpn1-mCherry-NatMX6 Nxt3-GFP-KanMX6 otr1R:ade6+ ade6-210 leu1-32 ura4-D18* | 4 |
| 9329 | *Cpn1-mCherry-NatMX6 Ubp3-GFP-KanMX6 otr1R:ade6+ ade6-210 leu1-32 ura4-D18* | 4 |
| 9413 | *sir2::ura4+ Cpn1-Flag-NatMX6 ade6-210 leu1-32 ura4-D18* | S4 |
| 9418 | *sir2::ura4+ cpn1::KanMX6 Mkt1-Flag-NatMX6 ade6-210 leu1-32 ura4-D18* | S4 |
| 9440 | *rik1::ura4+ Cpn1-GFP-KanMX6 Pabp-mRFP-HygMX6 leu1-32 ura4-D18* | 5 |
| 9520 | *h- ago1::ura4+ Pabp-GFP-KanMX6 otr1R:ade6+ ade6-210 leu1-32 ura4-D18* | 5,S5 |
| 9529 | *h- clr4::HygMX6:ura4+ Pabp-GFP-KanMX6 otr1R:ade6+ ade6-210 leu1-32 ura4-D18* | 5,6 |
| 9536 | *clr4::HygMX6 GFP-Cnp1-NatMX6 leu1-32 ura4-D18* | 6 |
| 9538 | *ago1::ura4+ cpn1::NatMX6 Pabp-GFP-KanMX6 ade6-210 ura4-D18 leu1-32* | 5,S5 |
| 9542 | *clr4::HygMX6:ura4+ cpn1::NatMX6 Pabp-GFP-KanMX6 ade6-210 leu1-32 ura4-D18* | 5 |
| 9596 | *clr4::HygMX6:ura4+ dhp1-2 (NatMX6) otr1R:ade6+ ade6-210 leu1-32 ura4-D18* | 6 |
| 9609 | *clr4::HygMX6:ura4+ cpn1::KanMX6 dhp1-2 (NatMX6) otr1R:ade6+ ade6-210 leu1-32 ura4-D18* | 6 |

**Table S2.**
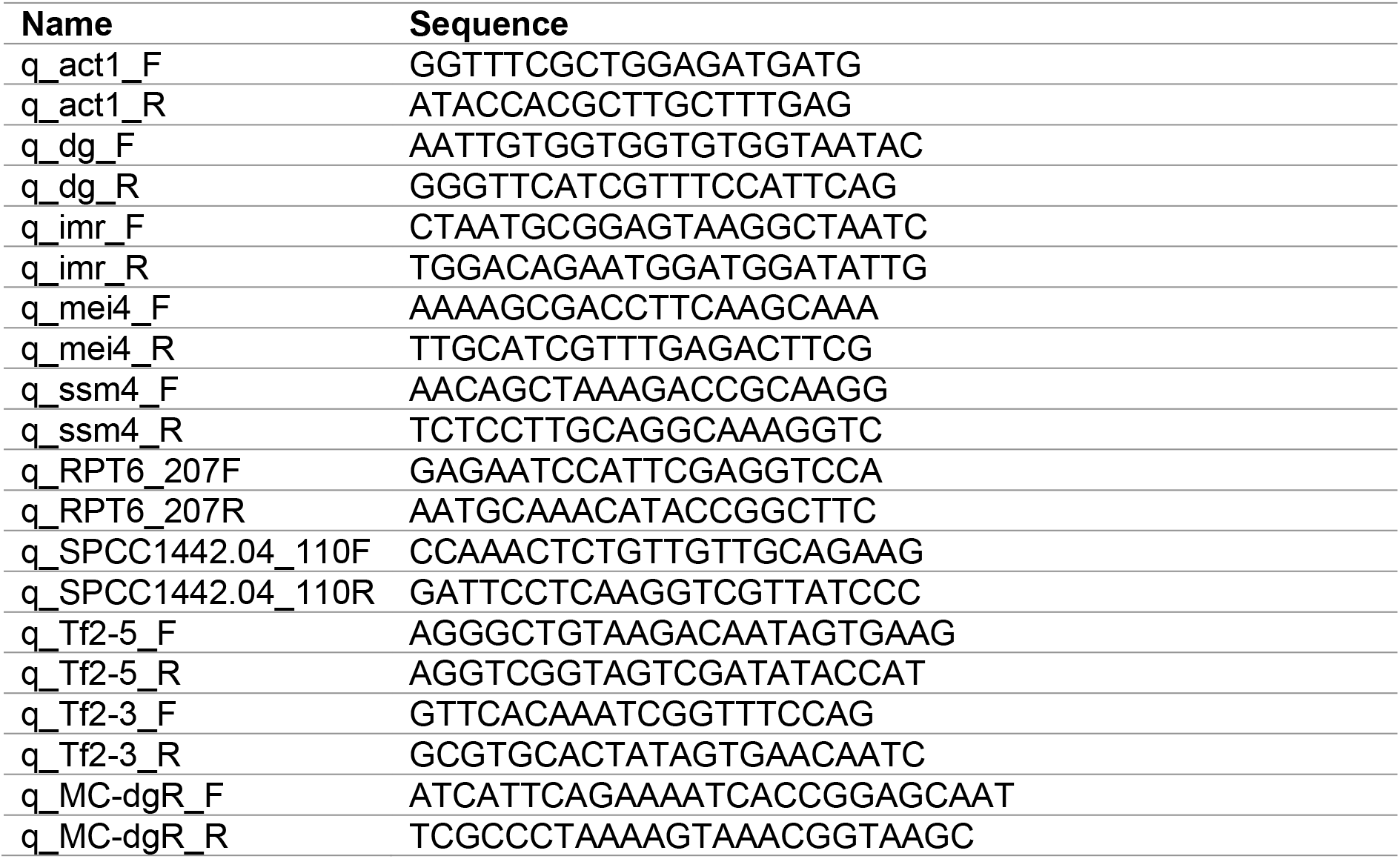
qPCR primers.

**Table S3.**
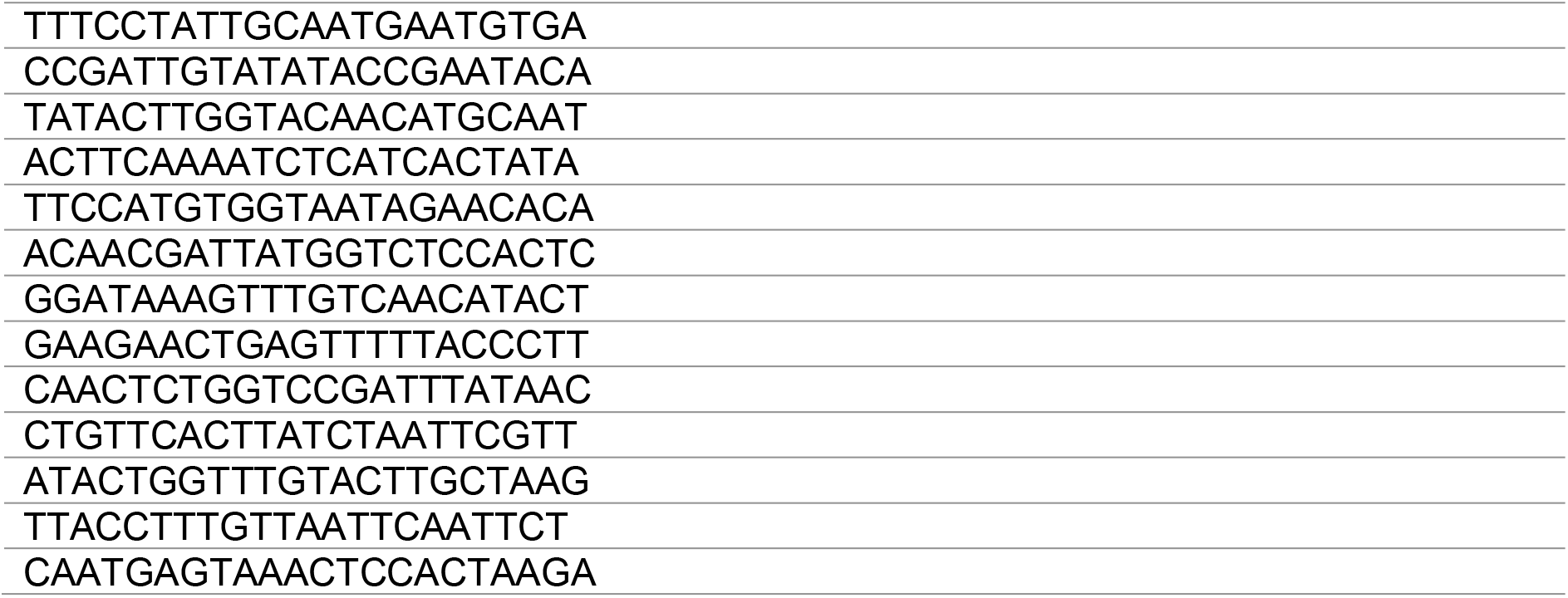
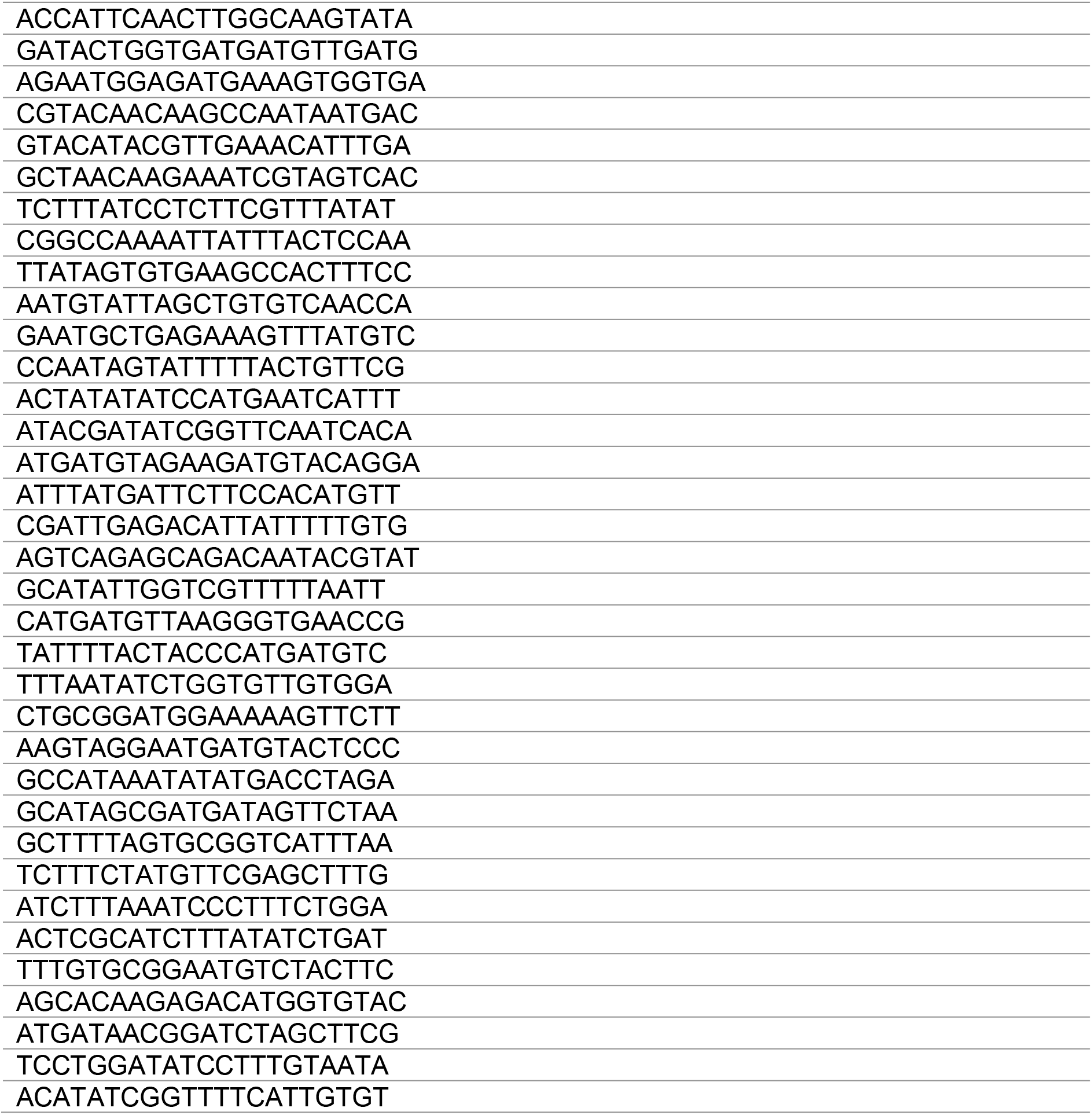
smRNA FISH probes.

